# The Ambiguous Genetic Code of Methanogenic Archaea that Grow on Methylamines

**DOI:** 10.1101/2025.06.11.659114

**Authors:** Katie E. Shalvarjian, Grayson L. Chadwick, Paloma I. Pérez, Philip H. Woods, Victoria J. Orphan, Dipti D. Nayak

## Abstract

Natural genetic code expansion is a phenomenon wherein an additional amino acid is encoded by a stop codon. These non-standard amino acids are beneficial as they facilitate novel biochemical reactions. However, code expansion leads to ambiguity at the recoded stop codon, which can either be read through or terminated. Pyrrolysine (Pyl) is encoded by the amber codon (TAG/UAG) and is widespread in archaea, where it is required for methylamine-mediated methanogenesis, an environmentally important metabolism. Mechanisms to conditionally suppress the amber stop codon for Pyl installation during protein synthesis have not been identified. Using the model methanogen, *Methanosarcina acetivorans,* we demonstrate that Pyl-encoding archaea maintain an ambiguous genetic code wherein UAG encodes dual meaning as stop and Pyl. Our data suggest that expression of Pyl biosynthesis and incorporation genes is tuned to the cellular demand for Pyl, which allows these archaea to navigate ambiguous stop decoding in response to environmental cues.

## Introduction

The genetic code is a core feature of life on Earth; through this triplet code, information stored in the four nucleotides of mRNA are translated into the twenty standard amino acids found in peptides/proteins. While initially proposed to be immutable^1^, it is now well established that the genetic code is flexible and subject to evolutionary pressure. At least 33 alternative genetic codes have been identified across the tree of life—in ciliates^2^, bacteria^3–5^, archaea^6,7^, fungi^8^, and phage^9^—where either a sense or a stop codon is reassigned to deviate from the standard genetic code. Natural genetic code expansion is a distinct case of codon reassignment, wherein a stop codon is recoded to an amino acid beyond the standard twenty amino acids^10^. Natural genetic code expansion is beneficial because the non-standard amino acids can enhance reaction rates or facilitate novel biochemistry^11–14^. But, it also presents a unique challenge due to the ambiguous interpretation of the recoded stop codon: the ribosome can either read through the recoded stop codon to add the non-standard amino acid to the peptide chain or terminate translation by recruiting a release factor instead. This ambiguity likely leads to the production of two pools of proteins— an extended form and a truncated form—for all transcripts containing the recoded stop codon. In the case of Selenocysteine (Sec), the 21^st^ amino acid, a Sec-tRNA^Sec^ specific elongation factor called SelB directs the ribosome to incorporate Sec at a UGA codon contingent on the presence of a proximal structural motif called SECIS (selenocysteine insertion sequence)^15,16^. In contrast, neither a sequence motif nor a specialized elongation factor has been found for Pyrrolysine (Pyl), the 22^nd^ proteinogenic amino acid^17^. Thus, the molecular mechanisms that Pyl-encoding microorganisms use (if at all) to tolerate ambiguous stop decoding remains unclear.

Pyl is biosynthesized from two molecules of lysine^18^ and is installed in elongating polypeptides using a dedicated UAG-decoding tRNA^Pyl^ that is charged by a cognate tRNA synthetase (PylRS)^19^. Pyl was first identified in the active site of methylamine-specific methyltransferases in methanogenic archaea (methanogens)^13^, where it is hypothesized to play an essential role during the methyltransfer reaction from methylamines to a corrinoid-containing protein^14^. Since then, Pyl has been identified across both archaea^20–23^ and bacteria^24,25^. However, a mechanism for targeted insertion of Pyl is not known in any of these microorganisms.

Here, we address the conundrum of how Pyl-encoding archaea have evolved to navigate a potentially ambiguous genetic code. First, we use an *in silico* approach to assess the distribution of Pyl across archaea and identify features that are unique to Pyl^+^ genomes. Although Pyl-encoding archaea have lower TAG stop codon usage overall, it can vary substantially from 1 to 27%, in line with previous reports^7^. Hence, the pattern of stop codon usage is neither a universal solution for the ambiguous genetic code created by Pyl nor is it a reliable predictor for its presence. Next, we engineer a heterologous expression system in *Methanosarcina acetivorans*, a native Pyl user, to test the functionality of Pyl systems from metabolically and taxonomically diverse archaea *in vivo*. Our functional assays add a layer of nuance to computational predictions and demonstrate that the presence of the *pyl* genes alone is not sufficient evidence for genetic code expansion. Finally, we provide conclusive evidence that *M. acetivorans* has an ambiguous genetic code in which the amber codon can be interpreted either as a stop codon or as Pyl. To this end, we describe possible mechanisms through which *M. acetivorans* maintains an ambiguous genetic code without a detrimental impact to organismal fitness.

## Results

### Pyrrolysine is present in archaea that perform anaerobic methylamine and methane metabolism

To map the distribution of Pyl in archaea, we searched for genes involved in its biosynthesis and incorporation in reference genomes available in the Genome Taxonomy Database (n=4,416; GTDB 214.0)^26^. The biosynthesis genes for Pyl (*pylBCD*)^18^ are typically encoded in an operon^27,28^ and are generally found in close proximity to the incorporation genes, comprised of tRNA^Pyl^ (*pylT*)^29^ and its cognate aminoacyl tRNA synthetase (*pylRS*, denoted as *pylS* hereafter)^19,20,28^ (**Fig 1**). We observe a strong correlation between the presence of the biosynthesis (*pylBCD*) and incorporation (*pylTS*) modules (*p=*1.4e-273, χ^2^*=*1249, **Table S1**); genomes encoding both sets of genes were annotated as Pyl^+^ (putative Pyl users) and the remainder as Pyl^-^. Consistent with previous reports^7^, the Pyl^+^ genomes are broadly distributed across archaea (**Fig S1, Table S2**) but can be discontinuous at the Genus level. For instance, among the *Asgardarchaeota*, only a few strains within the Genus *Borrarchaeum* are Pyl^+^ (**Fig S1, S2A**)^7,21^. Likewise, even though 92% of *Methanosarcina* genomes are Pyl^+^ (**Table S3**), a few strains have lost Pyl (**Fig S1, S2B**).

**Figure 1.**
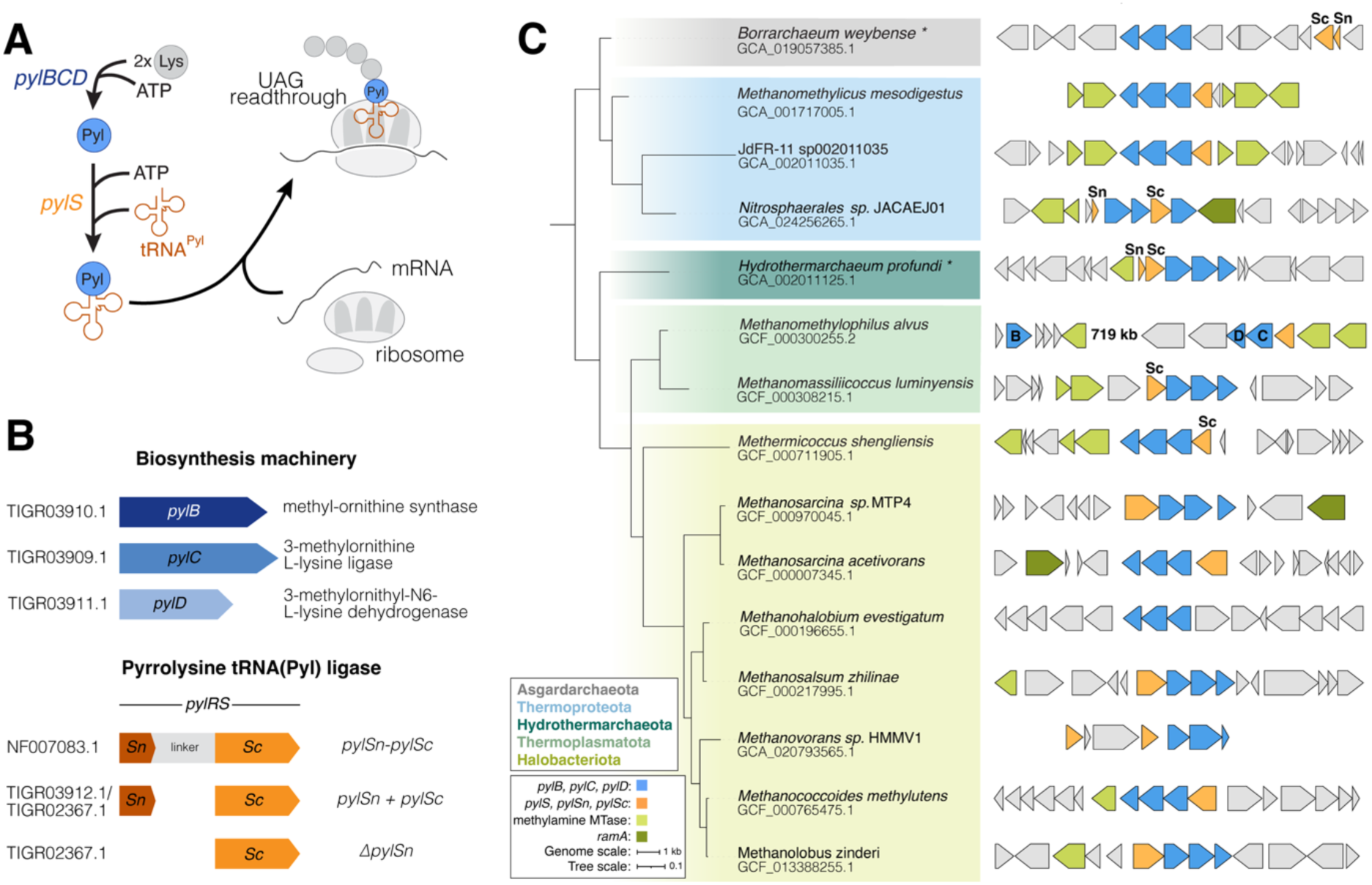
Distribution of Pyrrolysine (Pyl) in archaea. **A)** Proposed mechanism for Pyl biosynthesis (blue), and incorporation (orange) into elongating polypeptides by a Pyl-tRNA^Pyl^ charged by its cognate aminoacyl tRNA synthetase (PylS). **B)** Pyl^+^ archaea were annotated based on the presence of the biosynthesis genes encoded by *pylBCD* (in blue) and the Pyl aminoacyl tRNA synthetase (PylS; in orange). Genes and their corresponding Hidden Markov Models (HMM’s) accessions used for the bioinformatic survey are shown. Domain architecture and class distinctions are noted for each of the *pylS* classes. **C)** Pyrrolysine biosynthesis (blue) and incorporation (orange) genes and their genome neighborhood (±7.5 kb of *pylB,* unless otherwise noted) across different Phyla of Pyl^+^ archaea. The genome accession number is provided below the strain names. The corrinoid reductive activase *ramA* (dark green) and methylamine methyltransferases (light green) that often co-occur with the *pyl* genes are indicated. Strains denoted with an asterisk lack genes involved in methane metabolism. **Pyl:** pyrrolysine; **MTase**: methyltransferase; ***pylB***: methyl-ornithine synthase; ***pylC***: 3-methylornithine L-lysine ligase; ***pylD***: 3-methylornithyl-N6-L-lysine dehydrogenase; ***pylS***: pyrrolysine tRNA(pyl) ligase; ***ramA***: corrinoid protein reductive activase.

The Pyl machinery (i.e. PylBCD and PylTS) strongly co-occurs with two markers of methane metabolism in archaea: methyl-coenzyme M reductase (MCR) and component A2^30^ (*p*=3.1e-49, χ^2^=218, **Table S1**). If we survey only methanogens, the co-occurrence metric strengthens (*p=*1.1e-51, χ^2^=288 **Table S1**), a trend that does not hold true for **<u>an</u>**aerobic **<u>met</u>**hanotrophic archaea (ANME) (*p=*1.0, Fisher’s exact test). Pyl also co-occurs with genes encoding methylamine methyltransferases (*p=* 2.9e-231, χ^2^=1054, **Table S1**), although some Pyl^+^ archaea with these methyltransferases lack markers for methanogenesis (**Fig S2**) and are instead likely to perform acetogenesis^21^. Often, genes involved in methylamine metabolism^31^ are in close genomic proximity to the Pyl machinery (**Fig 1C, Fig S3**), which further corroborates their functional association.

*Methanosarcinaceae* from diverse environments commonly encode genes for methylamine-mediated methanogenesis and, thus, can be used to investigate fine-scale evolutionary events associated with Pyl-mediated genetic code expansion. To this end, we compared the gene trees for *pylS* and *pylBCD* to the 16S rRNA tree for this Family. While the *pylBCD* tree is in strong agreement with vertical inheritance (**Fig S4A**), the *pylS* tree diverges substantially (**Fig S4B, S5**). These data suggest that the Pyl biosynthesis and incorporation modules can have distinct evolutionary histories despite being functionally interlinked.

### Pyl-mediated genetic code expansion does not lead to universal patterns of stop codon compensation

Given that Pyl incorporation relies on read through of the TAG codon, we examined differences in stop codon usage between Pyl^+^ and Pyl^-^ genomes. As expected, TAG usage is significantly lower in Pyl^+^ genomes (median=5.0%) compared to Pyl^-^ genomes (median=22.0%) (Welch’s t-test, *p=*5.6e-47) (**Fig 2A**). However, taxon-specific features, like GC content^32^, can be major confounders in these global trends. To control for these biases, we analyzed the distribution of stop codon usage and GC content within the *Methanosarcinaceae* and the *Methanomethylophilaceae*, two groups that have a relatively high distribution of Pyl^+^ and Pyl^-^ genomes. Despite a large difference in GC content, TAG usage in Pyl^+^ genomes from *Methanosarcinaceae* (median=4.9%) and *Methanomethylophilaceae* (median=4.1%) is similar (**Fig 2B**). Further, within *Methanosarcinaceae,* we observe a marked increase in TAG stop codon usage for Pyl^-^ strains to their Pyl^+^ counterparts (Pyl^-^: 13.5%; Pyl^+^: 7.8%) (**Fig 2C, Table S2**).

**Figure 2.**
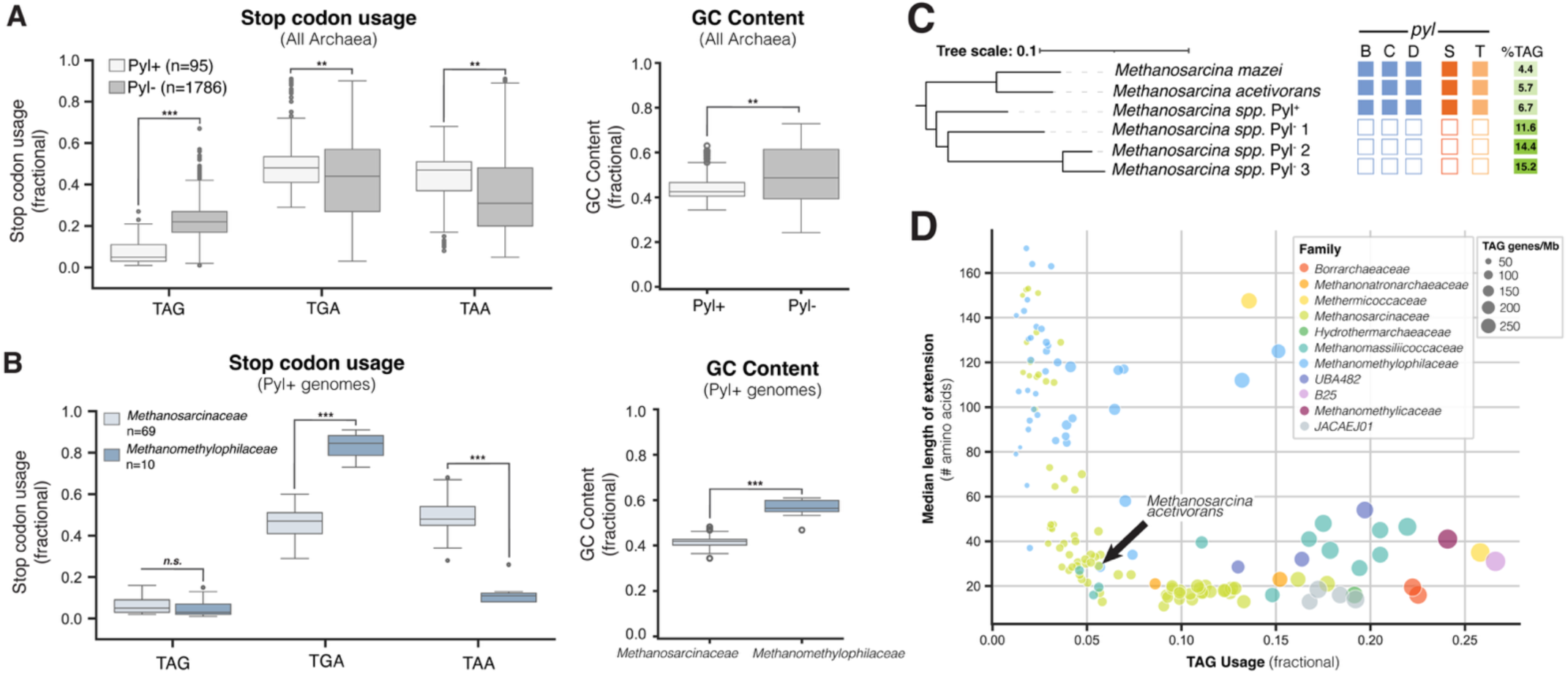
Stop codon usage in response to Pyl-mediated genetic code expansion. **A)** Box and whisker plots of stop codon usage and GC content in Pyl^+^ (light gray) and Pyl^-^ (dark gray) genomes within the archaea. **B)** Box and whisker plots of stop codon usage and GC content in Pyl^+^ genomes within the *Methanosarcinaceae* (light blue) and *Methanomethylophilaceae* (dark blue). Stop codon usage was calculated as fraction of total stop codons in protein encoding genes. All statistical tests for panels A and B were performed using Welch’s t-test (\*\*\**p*<1.0e-10, \*\**p*<1.0e-5, ***n.s.*** not significant). **C)** TAG usage in select *Methanosarcina* strains. The presence of a given gene is indicated by a filled-in square corresponding to the gene noted above the box; %TAG usage is indicated by the opacity of green box and denoted numerically. *Methanosarcina spp.* accessions are as follows: Pyl^+^: GCA_00171414685.2; Pyl^-^1: GCA_017883485.1; Pyl^-^2: GCA_003157235.1; Pyl^-^_3_: GCA_003164755.1 **D)** Genome-specific traits of Pyl^+^ archaea. A graph showing the relationship between the fractional TAG usage and the median extension length (# of amino acids) of putative Pyl-encoding proteins. The size of the genome dot indicates the relative abundance of putative Pyl containing genes (genes/Mb). The Family-level taxonomy is indicated by the color of the genome dot, and *Methanosarcina acetivorans* is indicated with the black arrow.

We did not detect a global pattern of compensation for decreased TAG usage in the Pyl^+^ genomes. While there are significant differences in TGA and TAA usage between Pyl^+^ and Pyl^-^ genomes across archaea (Welch’s t-test, *p=*8.3e-7 and *p*=5.9e-5, respectively) (**Fig 2A**), these are likely driven by taxon-specific factors. For instance, the Pyl^+^ genomes in *Methanomethylophilaceae* have higher TGA usage relative to *Methanosarcinaceae*, likely due to their heightened GC content (**Fig 2B)**. Hence, even though reduced TAG usage correlates with the presence of Pyl, there is no global compensation pattern. Rather, changes in TAA and TGA usage are contingent on factors like GC content.

Although TAG codon usage is substantially lower in Pyl^+^ genomes, it can be highly variable, from 1% to 27% (**Table S2**), in agreement with previous reports^7,22^. We hypothesized that differences in TAG codon usage might impact the fraction and function of putative Pyl-containing proteins Pyl^+^ genomes. Hence, we predicted ORFs with internal TAG codons and measured their putative readthrough lengths within each Pyl^+^ genome (**Table S4**). Overall, we observe that heightened TAG usage (>15%) is associated with an increase in the density of putative Pyl-containing proteins, whereas low TAG usage (<5%) is associated with an increase in readthrough length of TAG-containing ORFs (**Fig 2D, Fig S6**).

### Genetic studies of diverse Pyl systems defy computational predictions

Although Pyl is broadly distributed in archaea, its role in genetic code expansion has only been studied *in vivo* in the context of a few methanogens^7,33,34^. To broaden our functional understanding of Pyl across the domain, we expressed the Pyl machinery from two non-methanogens in the genetically tractable model organism, *Methanosarcina acetivorans.* The first strain, *Borrarchaeum weybense*, is an *Asgardarchaeon* hypothesized to use Pyl to perform methylamine-mediated acetogenesis^21^. The second strain, ANME-3 S7, contains Pyl despite lacking methylamine metabolism^22,35^. Axenic cultures are currently unavailable for both these strains, which prevents attempts to investigate their Pyl machinery in its native context. Moreover, these strains are unusual because they break from the strong co-occurrence patterns shown in (**Fig S1, Table S1**) and thus might offer additional insights into the role of Pyl in archaea.

To develop *M. acetivorans* as a heterologous host, we generated deletion mutants lacking either the biosynthetic operon (*ΔpylBCD*) or the incorporation genes (*ΔpylTS*) using our well-established CRISPR-Cas9 editing system^36^. Neither mutant had off-target effects or suppressor mutations based on whole genome resequencing (**Table S5**). We observed that both mutants have a mild growth defect (slower by 15%; **Fig 3A, Table S6**) when Pyl is not essential (i.e., on methanol) and are unable to grow when Pyl is essential [i.e., on trimethylamine (TMA)]. Complementation of each deletion mutant with the native locus *in trans* restored growth across all conditions (**Fig 3A, Fig S7, Table S6**), indicating that no extragenic effects stem from the chromosomal deletions. To validate the proposed catalytic role for Pyl in methylamine methyltransferases^13,14,28,37^, we generated a suite of non-synonymous point mutations at the Pyl residue in the TMA-specific methyltransferase, *mttB1.* We expressed these point mutants *in trans* from a tetracycline-inducible promoter^38^ in the *ΔpylBCD* background, where the chromosomal copy of *mttB1* would be prematurely truncated at the TAG codon (**Fig 3B)**. If Pyl (O) is catalytically essential, then the mutant alleles of *mttB1* will not be able to rescue growth on TMA (**Fig 3A, Table S6**). Indeed, neither the *mttB1*_O334A_ nor the *mttB1*_O334K_ mutants restored growth on TMA (**Fig 3C, Fig S7, Table S6**). To control for polar effects caused by premature translational termination of chromosomal *mttB1,* we jointly expressed *mttB1*_O334K_ with its cognate corrinoid containing protein, *mttC1*, and observed the same lethal phenotype (**Fig 3C, Fig S7, Table S6**). Together these data affirm the hypothesis that Pyl is essential for the activity of methylamine methyltransferases, which underlies the lethal phenotype for the Δ*pyl* mutants on TMA.

**Figure 3.**
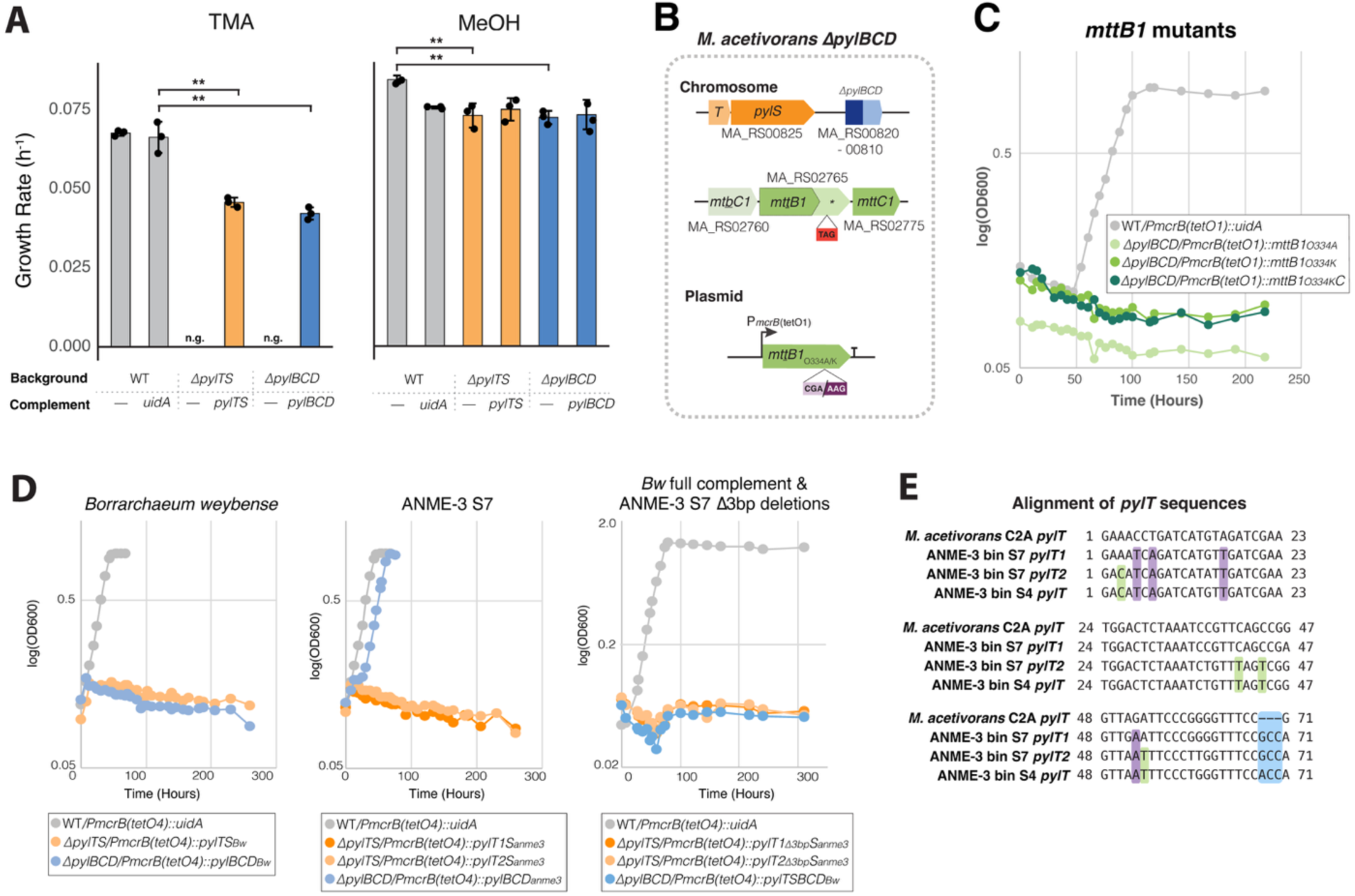
Heterologous expression of *pyl* genes from uncultivated archaea in *Methanosarcina acetivorans*. **A)** Growth rates of the parent strain (WWM60, denoted as wildtype or WT; in gray), the Δ*pylTS* (in orange), Δ*pylBCD* (in blue) mutants and the corresponding complementation strains in minimal media with either trimethylamine (TMA) or methanol (MeOH). In the WT strain, a control gene (*uidA)* was expressed from the tetracycline inducible promoter (P*mcrB*(*tetO1*)) on a plasmid like the genes complemented in *trans.* A two-sided t-test with unequal variances was used to compare mean growth rate from three biological replicates (\*\**p*<0.005). Error bars represent the standard deviation of three replicates. **B)** Diagram representing the genomic background for *mttB1* point mutation experiments. In the *ΔpylBCD* background, the chromosomal copy of *mttB1* is maintained, but TAG is obligately read as a stop codon due to the absence of Pyl biosynthesis machinery. A point mutated form of *mttB1* is complemented *in trans* under the control of the tetracycline inducible promoter, P*mcrB*(*tetO1*). **C)** Representative growth of triplicate curves for WT with a control gene (*uidA*; in gray) or the Δ*pylBCD* strain with the *mttB1* point mutants (in green) after transferring MeOH-grown cells into minimal medium with TMA. **D)** Representative growth of triplicate curves for the *pyl* genes from *Borrarchaeaum weybense* or ANME-3 bin S7 (including Δ3bp deletion in *pylT).* Complements were expressed *in trans* under a tetracycline-inducible promoter (P*mcrB*(*tetO4*)) in a *M. acetivorans* Δ*pylTS* (orange) or Δ*pylBCD* (blue) mutant as indicated. All growth curves were performed by transferring MeOH-grown cells into medium with TMA. WT with a control gene (*uidA*) are shown in gray. **E)** Alignment of *pylT* sequences from two members of ANME-3 (bin S7 and bin S3) with *M. acetivorans*. Blue box highlights the 3 bp insertion identified and deleted in panel D. Purple box highlights point mutations shared amongst all ANME *pylT* sequences. Green box highlights point mutations shared between S7 *pylT2* and S3 *pylT.* In panels A, C, and D, all strains bearing a plasmid for expression were cultivated with puromycin to maintain the plasmid and 100 µg/ml tetracycline to induce full expression of the complement from the tetracycline-inducible promoter.

Next, to test the functionality of non-native Pyl machinery, we complemented each deletion mutant with the *pyl* genes from *B. weybense* or ANME-3 S7 under the control of a tetracycline-inducible promoter. Even when fully induced, *pylTS*, *pylBCD* or *pylTS+pylBCD* from *B. weybense* were unable to restore growth on TMA (**Fig 3D**). Given that *pyl* genes from archaea have been successfully expressed in bacteria^39^, these data suggest that the *B. weybense* Pyl system is likely non-functional. In contrast, *pylBCD* from ANME-3 S7 could restore growth of the Δ*pylBCD* mutant on TMA (**Fig 3D, Table S6**). We tested both copies of the *pylT* locus (*pylT1* and *pylT2*) from ANME-3 S7 with their cognate synthetase, and neither could support growth of the *ΔpylTS* mutant on TMA (**Fig 3D**). In an alignment of ANME-3 S7 *pylT* sequences with *M. acetivorans pylT,* we identified a 3bp insertion at the 3’ end of the tRNA that could alter its secondary structure and disrupt its function (**Fig 3E**). To test this hypothesis, we generated Δ3bp deletions of *pylT1* and *pylT2* from ANME-3 S7 (**Fig 3D**) and introduced it with its cognate synthetase in *M. acetivorans*. However, we were still unable to rescue growth on TMA. Altogether, our heterologous expression platform suggests that only the Pyl biosynthetic machinery derived from ANME-3 S7 is functional in *M. acetivorans*. However, despite retaining the capacity to produce Pyl, it seems unlikely that ANME-3 S7 can incorporate it into proteins.

### The amber codon can be read as a stop or as a Pyl residue in *Methanosarcina acetivorans*

Although Pyl^+^ genomes have lower TAG usage, it is not clear if this codon has been obligately recoded to Pyl. Our genomic analyses show that *M. acetivorans* exists at an inflection point on the spectrum of Pyl^+^ genomes (**Fig 2D**), which uniquely positions it as a model system to experimentally test for TAG recoding. To this end, we evaluated readthrough versus translational termination at the TAG codon in RdmS, a non-essential Pyl-containing sensor kinase^34,40^. We fused a 1xFLAG tag at the N-terminus of *rdmS* and expressed it *in trans* in either a *ΔrdmS* or a *ΔpylBCD* background. In the latter, RdmS would be obligately truncated at the TAG codon located in its second PAS domain (**Fig 4A**). As expected, we only detected this “short-form” protein of approximately 25 kDa by α-FLAG western blot in the *ΔpylBCD* mutant (**Fig 4B, Fig S8**). In contrast, both the “long -form” and “short -form” of FLAG-tagged RdmS were detected in the Pyl^+^ *ΔrdmS* background (**Fig 4B, Fig S8**). Since a UAG codon retains the ability to recruit a release factor for translation termination, even in the presence of Pyl-tRNA^Pyl^, we searched for genes that might use it as a bona fide stop codon in *M. acetivorans* (see Methods). One candidate is *rnfA*, a membrane integral subunit of the energy conservation complex RNF^41,42^. If TAG is read as Pyl in lieu of a stop, RnfA would have a putative 28 AA C-terminal extension. Yet, *rnfA* homologs across *Methanosarcinaceae* lack the 28 AA C-terminal extension and use either TAA or TGA as a stop codon instead (**Fig S9, Table S7**), supporting a stop interpretation for TAG in *rnfA* from *M. acetivorans*. Taken together, these data support the hypothesis that the amber codon encodes a dual meaning in *M. acetivorans*.

**Fig 4.**
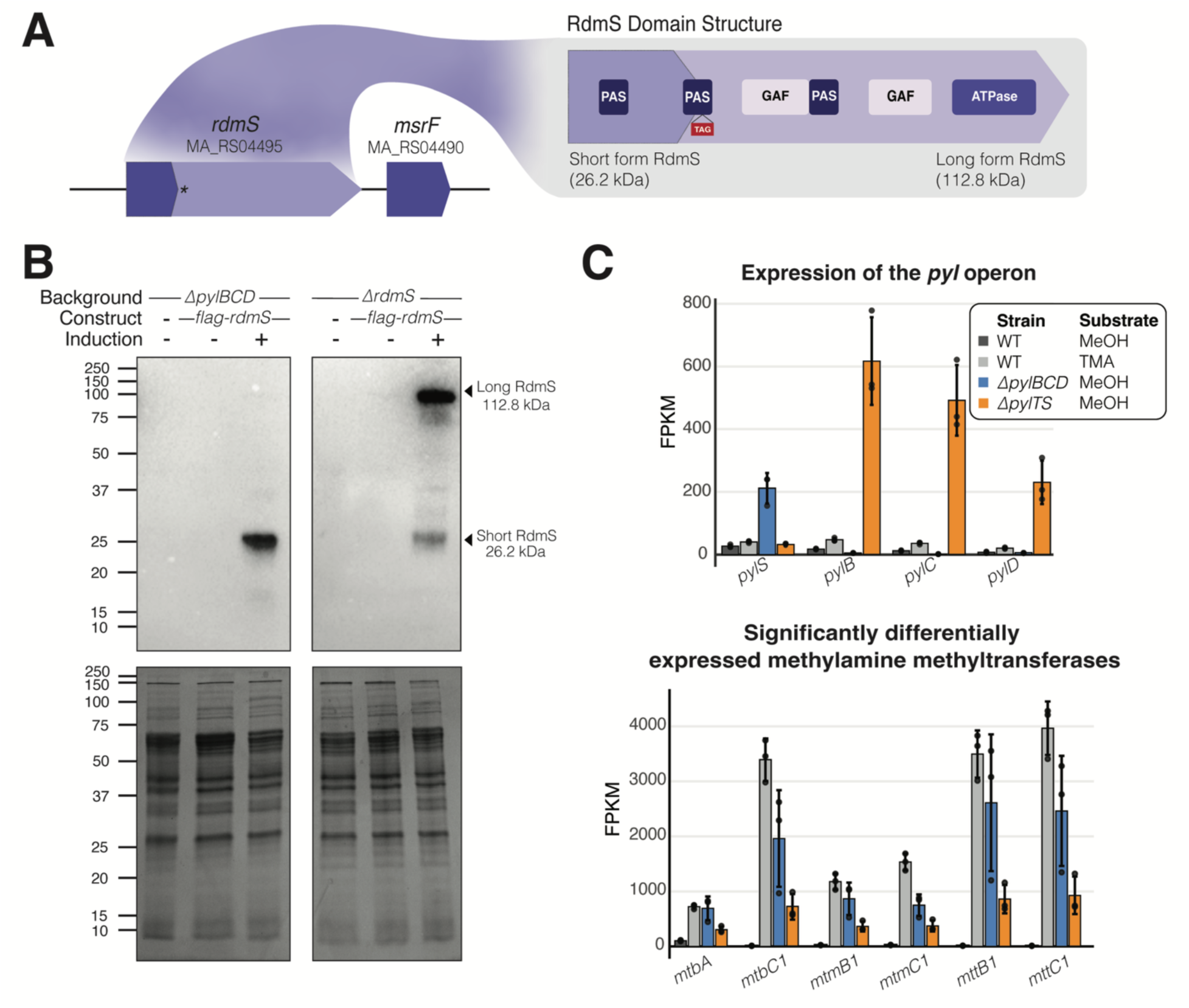
The amber codon can be interpreted as a stop codon or Pyl in *Methanosarcina acetivorans*. **A)** Diagram of RdmS domain structure and genomic organization. RdmS is a predicted sensor kinase located upstream of *msrF*^56^, a Msr (<u>m</u>ethanol-<u>s</u>pecific <u>r</u>egulator) family protein likely involved in the regulation of methylsulfide methyltransferases. Asterisk and TAG box indicate the location of the Pyl residue, where truncation would result in a ∼26.2 kDa protein. **PAS: P**er-**A**rnt- **S**im domain; **GAF:** c**G**MP-specific phosphodiesterases, **A**denylyl cyclases and **F**hlA; **ATPase**: **ATP** hydrolyzing domain. **B)** α-FLAG western blot for a N-terminal 1xFLAG translational fusion of *rdmS* expressed *in trans* from tetracycline inducible promoter, P*mcrB*(*tetO4*). Each blot represents a different genetic background, where induction of the 1xFLAG translational fusion of *rdmS* is indicated as no induction (-) at 0 µg/ml tetracycline or maximum induction (+) at 100 µg/ml tetracycline. Predicted molecular weights for the long form RdmS protein (where Pyl is incorporated at TAG) and the short form RdmS protein (where TAG is read as a stop) are indicated on the right side of the gel. Coomassie stained SDS-PAGE gels shown beneath western blot as loading control. **C)** Fragment per kilobase per million reads (FPKM) values of the *pyl* genes (top) and methylamine methyltransferase genes (bottom) in the Δ*pylBCD* mutant on methanol (MeOH) (blue), Δ*pylTS* mutant on MeOH (orange), WWM60 (parent strain also referred to as wildtype or WT) on MeOH (light gray) or trimethylamine (TMA) (dark gray). Error bars represent the standard deviation amongst plotted FPKM values for three replicates. The DESeq2 data can be found in Table S8.

In the absence of any sequence-specific cues for conditional suppression of the amber codon, we hypothesized that *M. acetivorans* might use the environmental context to dynamically tune the expression of the *pyl* genes i.e. produce more *pyl* transcripts in the presence of methylamines. Indeed, when we compare the global transcriptome, we observe that all *pyl* genes are substantially upregulated on TMA compared to methanol (**Fig 4C**, **Table S8**). Interestingly, even in the absence of TMA, *pylTS* is upregulated in the *ΔpylBCD* mutant and vice versa (**Fig 4C, Table S8**). These transcriptional profiles suggest that the *pyl* genes are dynamically expressed based on the cellular demand for Pyl in both a substrate-dependent and substrate-independent manner.

### Archaeal release factor copy number is not correlated with Pyl

Besides the expression of the *pyl* genes, the abundance and/or specificity of the archaeal release factor, aRF1, could, in principle, bias translation termination of the amber codon. A previous study showed that *Methanosarcina barkeri* has two copies of aRF1 and that one of them recognizes all three stop codons *in vitro*^43^. *M. acetivorans* also encodes two copies of aRF1, aRF1-1 (MA_RS00215) and aRF1-2 (MA_RS05270), but their functions have not been investigated. Thus, we reasoned that Pyl^+^ archaea might encode more than one aRF1, perhaps with distinct affinities for stop codons and expression levels, which could bias how UAG is decoded. We mapped the distribution of aRFs across all archaea and found that 4.3% of sampled genomes encode two copies of aRF1 (n=81, **Table S9**). However, most Pyl^+^ genomes with two copies of aRF1 belong to the Genus *Methanosarcina* (**Table S9**). Thus, it is likely that the additional aRF1 copy in *Methanosarcina* spp. arose from gene duplication due to lineage-specific genome expansion. Accordingly, we found that aRF1 copy number increases with genome size in archaea (*p=*2e-21, **Fig S10A**), independent of predicted Pyl use (**Fig S10B**). Finally, our transcriptomics analyses indicate that the aRF1 genes are constitutively expressed in *M. acetivorans* across all tested conditions (**Fig S10C**). While our data do not rule out the possibility for post-transcriptional regulation or differing affinities of aRF1s, it seems unlikely that aRF1 duplications play a universal role in streamlining an ambiguous code in Pyl^+^ archaea.

## Discussion

Pyl-based code expansion is a double-edged sword: the Pyl residue enables methylamine-metabolism but also leads to ambiguous interpretation of the amber codon. Here, we outline two co-occurring processes that allow Pyl^+^ archaea to maintain Pyl in their proteome. First, the acquisition of *pyl* genes selects for the genome-wide redistribution of stop codons to lower TAG usage (**Fig 2**). This process is relatively swift and can be observed in archaea that have recently gained Pyl, like *Hydrothermarchaeum profundi* (**Fig S1**), or in archaea that have recently lost Pyl, like ANME-3 S7, which encodes partially functional *pyl* genes (**Fig 3D**), or the *Methanosarcina spp.* that have entirely lost *pyl* genes (**Fig 2C**). Given the speed and ease at which this process occurs, lowering TAG usage is likely the first step in maintaining Pyl.

Once TAG usage stabilizes, the ability to tolerate ambiguous decoding at all remaining amber codons is essential to maintaining Pyl homeostasis. Using RdmS, a gene with an in-frame TAG codon, we provide evidence that, at least in *M. acetivorans,* the amber codon can be simultaneously read as both a sense and nonsense codon *in vivo* (**Fig 4**). These data reinforce previous findings that show the co-existence of the extended and truncated forms of the Pyl-containing monomethylamine-methyltransferases (MtmB) in *Methanosarcina barkeri* MS^33^. Hence, unlike Sec, an ambiguous code is not an evolutionary intermediate but an endpoint in Pyl-mediated genetic code expansion.

To address how an ambiguous code might be tolerated, we present a heuristic model for *M. acetivorans* using our transcriptome data. In the absence of a sequence-specific suppression mechanism, we assume that the likelihood of readthrough and truncation are equal for all UAG-containing transcripts in the cell. Hence, we can define readthrough efficiency for a given UAG-containing transcripts as the ratio of the readthrough rate to the truncation rate, which is proportional to the ratio of Pyl supply and demand in the cell, respectively (see Methods). We can estimate Pyl supply by measuring the abundance of transcripts for the *pyl* genes, and Pyl demand by measuring the total number of UAG-containing transcripts (Eq. 1, see methods, FPKM:

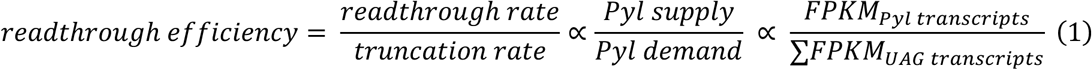

Based on equation 1, the readthrough efficiency is *ca.* two-times higher on methanol than on TMA i.e. there is a higher fraction of the long-form of the protein for all UAG-containing transcripts on methanol than on TMA (**Table S10**). At first glance, this outcome might seem counterintuitive, as the cells needs to produce Pyl-containing methylamine methyltransferases on TMA. However, a 4.0-fold increase in expression of the *pyl* genes (i.e. Pyl supply) is far surpassed by the dramatic increase in UAG-containing transcripts (i.e. Pyl demand) on TMA, *ca.* 188.0-fold, primarily due to the upregulation of methylamine methyltransferases (**Fig 4C, Table S8**). Hence, even though Pyl supply increases during growth on TMA, its demand far outpaces its supply, leading to lower readthrough efficiency (**Fig. 5**).

**Fig 5.**
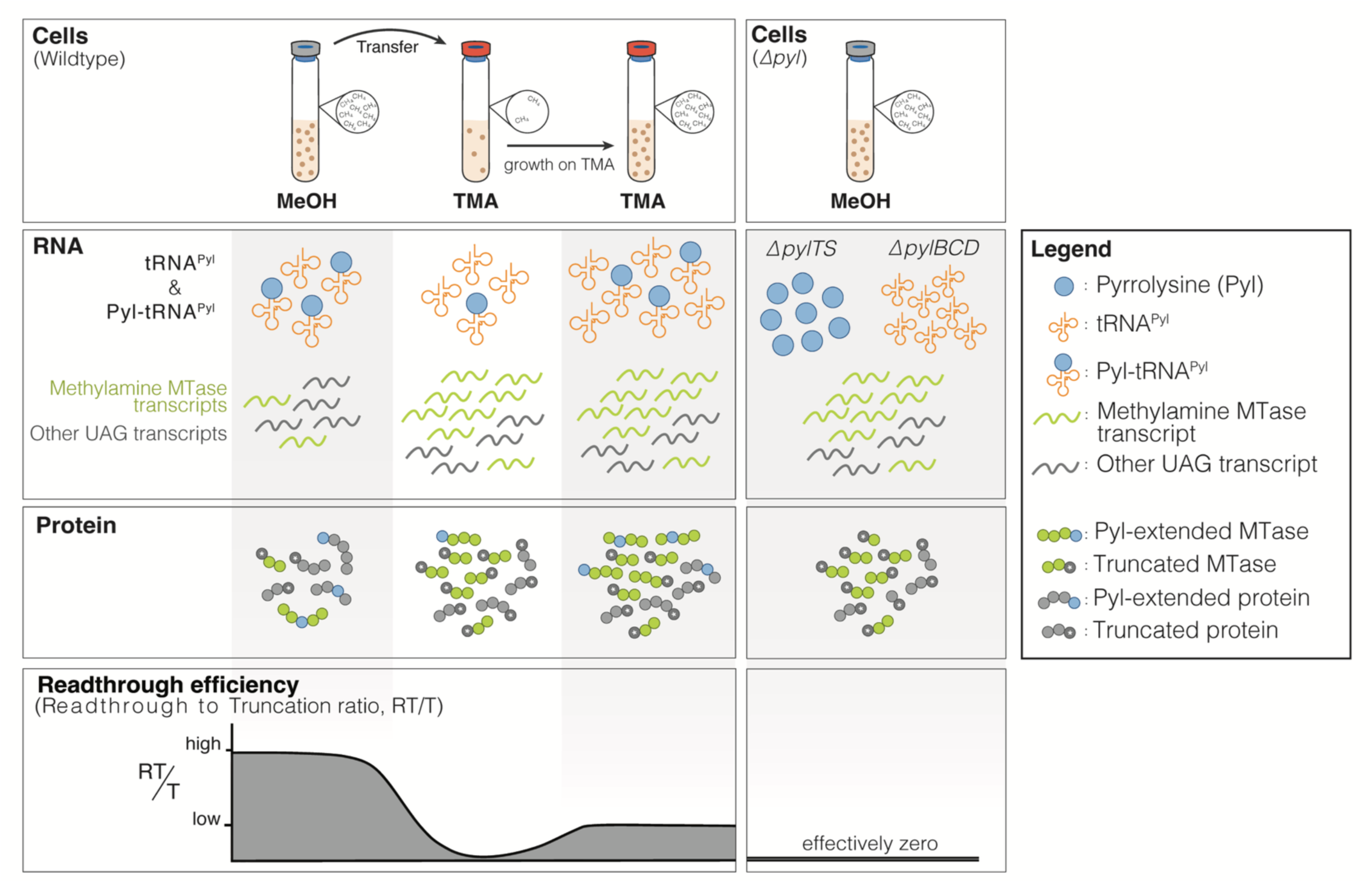
A model for Pyrrolysine (Pyl) homeostasis in *Methanosarcina acetivorans*. During growth on methanol (MeOH), the pool of UAG-containing transcripts is lower, leading to the higher readthrough efficiency (RT/T, bottom panel). We hypothesize that as cells switch from MeOH to trimethylamine (TMA), they encounter a transient state wherein the expression of methylamine methyltransferases (MTase) increases rapidly and Pyl demand outpaces its production. As a result, the readthrough efficiency transiently decreases. This decrease in RT/T likely leads to an increase in expression of *pyl* genes, and methylamine MTases, however, because the MTases have an outsized contribution to the total UAG transcript pool, MTase transcripts effectively outcompete other UAG transcripts for Pyl. We suggest our *pyl* knockouts have a readthrough efficiency that is close to zero and that they respond to this by upregulating the *pyl* genes as well as the methylamine-specific MTases as seen in **Fig 4C**.

In our *ΔpylBCD* and *ΔpylTS* knockouts, we eliminate Pyl supply and effectively reduce the readthrough efficiency to zero. Under this regime, we observe that *pylBCD* is upregulated in the absence of *pylTS*, and vice versa, on methanol (**Fig 3C**). While the cue(s) leading to an upregulation of the complementary *pyl* genes in the knockout mutants is unknown, we rule out sensing of free Pyl, as the transcriptomes of *ΔpylBCD* and *ΔpylTS* are nearly overlapping (**Table S8**). Instead, we hypothesize that these transcriptional changes stem from an inability to incorporate Pyl in a UAG-containing transcript, i.e. by sensing a diminishingly low readthrough efficiency. This artificially low readthrough efficiency, and its resulting transcriptional response, is likely akin to what occurs in wildtype cells upon initial exposure to TMA. Here, the rapid increase in demand for Pyl (due to expression of methylamine methyltransferases) results in a transiently low readthrough efficiency, which is partially recovered by increased expression in *pyl* genes (**Fig 5**).

Given the enrichment of TAG codons in genes with proposed regulatory function, one hypothesis for how cells sense readthrough efficiency is through a Pyl-containing repressor that is active when there is readthrough and inactive in its truncated form. In this scenario, high readthrough efficiency on methanol maintains the repressor in its elongated form and partially represses *pyl* genes. In contrast, during growth on TMA, the repressor is prematurely truncated at its UAG site, which results in de-repression of *pyl* genes (**Fig 5**). Therefore, when we impose a readthrough efficiency of zero through deletion of either *pylTS* or *pylBCD,* the repressor is fully inactive. This leads to a TMA-like transcriptional response on methanol, complete with upregulation of the remaining *pyl* genes and methylamine methyltransferases (**Fig 4C, Table S8**). Alternately, other mechanisms like attenuation or transcript stability may play a role in regulation too. It is worth noting that *pylB* has an in-frame TAG codon at the C-terminus. While the functional consequence (if at all) of Pyl in PylB^34^ is unknown, it is tempting to hypothesize that it plays a role in tuning Pyl biosynthesis.

Our model also explains the counterintuitive growth data wherein overexpression of the *pyl* genes is tolerated on methanol but leads to a growth defect on TMA (**Fig 3A**). An increase in Pyl supply likely elevates the readthrough efficiency across all growth conditions. This is tolerated on methanol since the readthrough efficiency is already high. In contrast, on TMA, where the readthrough efficiency is low, elevated Pyl incorporation in some UAG-containing transcripts might lead to a buildup of the “long form” of certain proteins that are detrimental to growth.

Altogether, we propose that Pyl^+^ archaea, like *M. acetivorans,* are expert funambulists that navigate the tightrope of Pyl homeostasis through transcriptional regulation of Pyl supply and demand. While we cannot rule out unknown mechanisms for sequence-specific Pyl insertion, we show that the ambiguity imposed by its presence can be tolerated successfully.

## Acknowledgements

We thank all members of the Nayak lab for their valuable feedback and support. DDN acknowledges funding from the Searle Scholars Program sponsored by the Kinship Foundation, the Rose Hills Innovator Grant, the Beckman Young Investigator Award sponsored by the Arnold and Mabel Beckman Foundation, the Alfred P. Sloan Research Fellowship sponsored by the Sloan Foundation, the Simons Foundation Early Career Investigator in Marine Microbial Ecology and Evolution Award, and the Packard Fellowship in Science and Engineering sponsored by the David and Lucille Packard Foundation. DDN is a Chan-Zuckerberg Biohub – San Francisco Investigator. KES was supported in part by the NSF Graduate Research Fellowship Program (Fellow ID: 202299857). GLC was supported by the Miller Institute for Basic Research in Science, University of California Berkeley. PIP was supported by University of California Berkeley’s Molecular and Cell Biology (MCB) department and the National Institute of Health (NIH) Bridges to Baccalaureate Program. DDN, PHW, VJO acknowledge funding from the Department of Energy through project number S589706. The funders had no role in the conceptualization and writing of this manuscript or the decision to submit the work for publication.

## Author contributions

KS was involved in conceptualization, data curation, formal analysis, methodology, investigation, validation, visualization, writing – original draft preparation, and writing – review and editing. GLC was involved in conceptualization, methodology, investigation, writing – original draft preparation, and writing – review and editing.

PIP was involved in conceptualization, methodology.

PHW and VJO provided resources.

DDN was involved in conceptualization, data curation, funding acquisition, investigation, methodology, project administration, resources, supervision, validation, writing – original draft preparation, and writing – review and editing.

## Data availability statement

Sequencing data have been deposited in the Sequencing Reads Archive and the BioProject number will be made available upon publication. All other data generated in this study are provided in the manuscript.

## Declaration of interests

The authors declare no competing interests.

## Methods

### Bioinformatic screen for Pyl and Pyl-associated genes

All archaeal genomes from GTDB Release 214.0^44^ were annotated with Prokka^45^ v.1.14.6 using the prokaryotic genetic code 11 to identify homologs of the Pyl biosynthetic genes (*pylBCD*), Pyl incorporation gene (*pylS*), methyl-coenzyme M reductase catalytic subunits (*mcrABG*), component A2, and methylamine-specific methyltransferases (*mttB, mtbB,* and *mtmB*). The Pyl-tRNA^Pyl^ sequences (*pylT*) were predicted in the Prokka pipeline using Aragorn^46^, and hits were counted for each genome using a custom script that can be found at https://github.com/kshalv/pyl. Gene identification was performed using a command line tool developed for automated gene searching with profile HMM’s (accessions can be found in **Table S12**) using the ‘hmm’ option, as previously described in [47] and can be found at: https://github.com/kshalv/hmm_tools. Pyl^+^ genomes were strictly defined as those containing a full suite of biosynthetic genes (*pylBCD*) and incorporation genes (*pylS*, *pylT*). In principle, a genome with just *pylTS* (0.2% of genomes) could be Pyl^+^ but there are no known Pyl transporters, so we also classified this group as Pyl^-^.

### Phylogenetic analyses

Phylogenetic trees depicted in **Fig 1C**, **Fig S1**, **Fig S2**, and **Fig S3** were generated using ete3 and a custom python script to parse the ar53_r214.tree available through Genome Taxonomy Database (GTDB). Genomes with a checkM ≥90% were used for all trees except for Fig S1 for which genomes with a checkM completeness scores of ≥80% were used. The CheckM score was calculated using a concatenated biomarker set described in [44]. The trees depicted in all figures, except for Fig S3-S5, were visualized using iTol v7.1^48^. Genomic neighborhood diagrams for genes +/- 7.5 kb up and downstream of *pylB* in Fig. S3 were generated using BioPython and overlaid on the GTDB tree using a custom script found at https://github.com/kshalv/pyl. Amino acid sequences used to generate gene trees for the Pyl biosynthetic and incorporation machinery within the Family *Methanosarcinaceae* were extracted from the corresponding GTDB 214.0 genomes (code 11) using the ‘seq’ option of the hmm_tools CLI described above with the ‘trusted cutoff’ threshold. Sequences were aligned using MAFFT and trimmed to 90% with TrimAl. The trimmed alignments were then concatenated using a custom script (https://github.com/kshalv/pyl). Gene trees were generated using the Geneious RAxML plug-in 8.2.11 and GAMMA BLOSUM62 substitution matrix with n_bootstraps=100. The 16S rRNA tree for the Family *Methanosarcinaceae* was generated using an alignment provided in GTDB r214.0 and assembled using the Geneious RAxML 8.2.11 plug-in. Tanglegrams were visualized using Dendroscope 3.5.9^49^. Sequences used to generate methyltransferase gene trees were extracted using the protocol described above from GTDB 214.0 genomes (annotated to code 15, with TAG recoding to Pyl using a custom script (https://github.com/kshalv/pyl). Sequences were dereplicated to 90% sequence identity using MMSeqs2^50^ easy-cluster. The tree was built using the Geneious RAxML 8.2.11 plug-in and was rooted to the methanol-specific methyltransferase family in iTol (v 7.1).

### Evaluation of stop codon usage and GC content

Stop codon usage was calculated for all genomes available in GTDB 214.0 using BioPython to extract the last three nucleotides of each annotated open reading frame (i.e., “CDS” designation) in the corresponding code 11 Genbank files generated by Prokka’s annotation pipeline. Usage for each stop codon was calculated as a fraction of the total number of predicted stop codons and was recorded for each genome. The GC content for each genome was derived from GTDB r214.0 metadata. Taxonomic groups were classified by binning accessions in accordance with the taxonomy designated by GTDB 214.0. All genomes used in this analysis exceeded a checkM genome completeness score of ≥90%. A Welch’s t-test was performed for each comparison group depicted in Figure 2 using the scipy.stats Python package.

### Identification of putative Pyl-containing proteins

Predictions for putative Pyl containing proteins were performed for each genome predicted to be Pyl^+^ based on the presence of 2/3 Pyl biosynthetic genes, the presence of *pylS,* and *pylT*. Each Pyl^+^ genome was reannotated using the Blepharisma Nuclear Code (code 15) in Prokka v1.14.6, effectively recoding each TAG codon to Glutamine (Gln, Q)^9^. For each genome, we identified putative Pyl proteins using a custom script, where CDS’s containing an in-frame ‘TAG’ codon were defined as putative Pyl proteins. For each Pyl protein, extension length was calculated as the difference between total length of the protein and the ‘TAG’ position and recorded in a genome-specific CSV file. We recorded the total number of putative Pyl proteins, the average and median extension lengths, and the density of putative Pyl proteins (i.e., normalized Pyl hits per Mb) for each genome. A CDS containing the TAG codon as putative stop codon was identified as a sequence where TAG readthrough results in a >5 AA extension for which there is 1) no predictable domain, and 2) is absent in sequence homologs.

### Media and culture conditions

All *Methanosarcina acetivorans* strains were grown without shaking at 37°C in bicarbonate-buffered high salt (HS) liquid medium containing either 50 mM trimethylamine hydrochloride (TMA) or 125 mM methanol as growth substrates. All growth substrates were added to the medium prior to sterilization, along with NH_4_Cl, Cysteine-HCl, and Na_2_S.9H_2_O to final concentrations of 19 mM, 2.8 mM, and 0.4 mM, respectively. Each strain was grown in a hermetically sealed Balch tube with N_2_/CO_2_ (80%/20%) in the headspace at 8-10 psi. All *Escherichia coli* strains were grown in lysogeny broth (LB) with antibiotic concentrations of 20 µg/mL chloramphenicol and/or 25 µg/mL kanamycin.

### Plasmid construction

Plasmids for Cas9-mediated genome editing were designed using the method previously described in [36]. PCR fragments containing the promoter (P*_mtaCB1_*), scaffolds, and terminator (T*_mtaCB1_*), were amplified with overhangs containing single guide RNAs (sgRNAs) that target Pyl genes (*ΔpylBCD,* MA_RS00820-MA_RS00810; *ΔpylTS* MA_RS00815). All sgRNAs were designed using the CRISPR site finder tool in Geneious Prime version 2023.2.1 and are provided in **Table S12**. PCR fragments containing the sgRNA scaffold were assembled into pDN201 linearized with *AscI* using the Gibson assembly method described previously in [51]. The repair template was assembled into the sgRNA containing derivative of pDN201 at the *PmeI* site. All complementation constructs were built using either pJK027A or pJK029A as described in **Table S12**. PCR fragments containing the genes of interest and if applicable, affinity-purification tags, were fused to the appropriate tetracycline-inducible promoter (**Table S12**) by linearizing pJK027A or pJK029A with *NdeI* and *HindIII*. Cointegrates of all plasmids with pAMG40 for expression in *M. acetivorans* were obtained using the Gateway BP Clonase II Enzyme Mix (Thermo Fisher, Waltham, MA, USA) and verified by restriction endonuclease digest. All cloning was performed in WWM4489^52^, a DH10B derivative, which was transformed by electroporation at 1.8 kV using an *E. coli* Gene Pulser (Bio-Rad, Hercules, USA), and the growth medium was supplemented with 10 mM Rhamnose to induce copy number as previously described [52]. All plasmids were extracted using Zyppy Miniprep Kit (Zymo Research, Tustin, CA, USA) and verified by Sanger sequencing at the UC Berkeley DNA Sequencing Facility. All primers and plasmids used in this study are listed in **Table S12** and **Table S13**, respectively.

### Methanosarcina acetivorans mutant generation

Liposome-mediated transformation was used to generate *M. acetivorans* mutants as previously described in [53]. Briefly, 20 mL cultures of *M. acetivorans* were grown in HS-Methanol (MeOH) medium until they reached mid-late exponential phase and were harvested for transformation. Transformants were plated and selected for on agar-solidified HS medium with 125 mM MeOH and 2 µg/mL puromycin. Plates were incubated in an intra-chamber anaerobic incubator maintained at 37°C with N_2_/CO_2_/H_2_S (1000 ppm/20%/balance) in the headspace. For all complementation strains, plasmids were screened by PCR and resulting products were verified by Sanger sequencing at the UC Berkeley DNA Sequencing Facility. For deletion mutants, individual colonies were picked and screened at the chromosomal locus using primers designated in **Table S12** for the desired mutation. Colonies that were positive for the desired mutation were then streaked on solid HS-MeOH medium with 20 µg/mL 8ADP (Carbosynth, San Diego, CA, USA) to cure the plasmid; single plasmid cured colonies were screened for the absence of the *pac* gene on the mutagenic plasmid with PCR. Chromosomal deletion mutants were sent for whole genome re-sequencing (described below) to screen for the deletion and suppressor mutations elsewhere on the chromosome. All strains used and generated in this study are listed in **Table S14**.

### DNA extraction and sequencing

Genomic DNA for whole genome re-sequencing of deletion strains was extracted from 10 mL of stationary phase culture of the *M. acetivorans* strain using the Qiagen blood and tissue kit following the manufacturer’s instructions (Qiagen, Hilden, Germany). Genomic DNA was sent to SeqCenter (Pittsburgh, PA) where it was further processed using the Illumina DNA Prep kit and sequenced (150 bp paired end reads). Sequencing reads were mapped to the *M. acetivorans* C2A reference genome using breseq^54^ version 0.35.5 using default parameters. All mutations identified through this analysis are listed in **Table S5**. Illumina sequencing reads for DDN121 (*ΔpylBCD*), and DDN146 (*ΔpylTS*) have been deposited in the Sequencing Reads Archive (SRA) and the project number will be made available upon publication.

### Cultivation for growth assays

All growth curves generated for this work were performed by pre-culturing strains in 10 mL HS-MeOH media at 37°C without shaking (HeraTherm General Protocol Microbiological Incubator, Thermo Fisher Scientific, Waltham, MA, USA) in hermetically sealed Balch tubes with N_2_/CO_2_ (80%/20%) in the headspace at 8-10 psi. All growth experiments were inoculated with 500 µl of the pre-culture at late exponential phase (OD_600_ ∼0.9-1.0). Growth of three independent biological replicates was measured by monitoring the optical density of cultures at 600 nm using a UV-Vis Spectrophotometer (Genesys 50, Thermo Fisher Scientific, Waltham, MA, USA) with an uninoculated “blank” containing 10 mL HS medium with the appropriate growth substrate. All strains harboring a plasmid for gene expression *in trans* also contained 2 µg/mL puromycin in the medium for plasmid maintenance and 100 µg/mL tetracycline for promoter induction. All cultures with tetracycline were wrapped in aluminum foil to prevent degradation over time. Growth rates were calculated as the slope of the linear fit of the log-transformed optical density versus time plot with a minimum five points with the highest R^2^ value (minimum cut-off ≥0.95).

### RNA extraction, sequencing, and analysis

10 mL cultures of WWM60 (parental strain), DDN121 (*ΔpylBCD*), and DDN146 (*ΔpylTS*) were grown in triplicate on HS + 125 mM MeOH at 37°C; an additional triplicate set of WWM60 was grown on HS+ 50 mM TMA at 37°C. When cultures reached mid-exponential phase (OD_600_∼ 0.45-0.65), 1 mL culture was removed for RNA extraction and mixed 1:1 with Trizol (Life Technologies, Carslbad, CA, USA) that was pre-warmed to 37°C. After a five-minute incubation at room temperature, 2 mL of 100% ethanol was added to each sample and RNA extraction was performed according to the manufacturer’s instructions using the Qiagen RNeasy Mini Kit (Qiagen, Hilden, Germany). RNA concentration was determined using Nanodrop One UV Spectrophotometer (Thermo Fisher Scientific, Waltham, MA, USA) before storage at −80°C. Samples were sent to SeqCenter (Pittsburg, PA) where each replicate underwent DNase treatment, rRNA depletion, library preparation, and Illumina paired-end sequencing (600 bp read length). Transcriptome analysis was performed using the KBase bioinformatics platform. Raw transcript reads (in FastQ format) were used as an input for read alignment to the *M. acetivorans* C2A reference genome using HISAT2 (v.2.1.0). The resulting read alignment was assembled using Cufflinks (v.2.2.1) and fold changes, significance values were calculated using DESeq2 (v.1.20.0). We considered changes in transcript abundance between pairwise comparison to be significantly differentially expressed if they exceeded a log_2_ fold change of ±2.0 and a *q*-value of ≤0.05. All DESeq2 data is recorded in **Table S7**. Raw transcript reads for DDN121 and DDN146 have been deposited in Sequencing Read Archive (SRA) and the project number will be made available upon publication; raw transcript reads for WWM60 grown on MeOH and on TMA have been deposited in the SRA.

### Immunoblotting of FLAG-tagged proteins

Immunoblotting was performed with cell lysates from a 10 mL culture in HS medium + 125 mM MeOH grown to late-exponential phase (OD600 ∼0.9-1.0). For strains harboring plasmids with tetracycline inducible genes, 2 µg/mL puromycin was added to maintain the plasmid and 100 µg/mL tetracycline was added to induce gene expression. Crude cell lysates were prepared from cultures using the method described in [51, 55]. Briefly, the culture was spun down at 5,000 x *g* at 4°C for 10 minutes using the Sorvall Legend XTR centrifuge (Thermo Fischer Scientific, Waltham, MA, USA). The culture supernatant was decanted, and the pellets were resuspended in 200 µL lysis buffer (50 mM Na_2_HPO_4_) and kept on ice for ten minutes prior to adding 1 µL of DNase along with 12 µl 5M NaCl. After mixing gently, the lysate was stored at room temperature for two minutes prior to centrifugation at 14,000 x *g* for ten minutes twice to obtain cleared lysate. Near-equal loading of 30 µg of protein was estimated using Bradford reagent (Sigma-Aldrich, St Louis, MO) with bovine serum albumin (BSA) standards per the manufacturer’s instructions. Cell lysates were run on a 12% mini-Protean TGX Gels (Bio-Rad, Hercules, USA) and transferred to a Trans-Blot Turbo Mini 0.2 µM PVDF membrane using the Bio-Rad Trans-Blot Turbo Transfer System (Bio-Rad, Hercules, USA). FLAG-tagged proteins were probed by immunoblotting using monoclonal anti-FLAG M2-Peroxidase (HRP) antibody (Sigma-Aldrich, St Louis, MO, USA) at a 1/50,000x dilution. Subsequent signal detection was performed using Immobilon Western Chemiluminescent HRP Substrate (Millipore, Burlington, MA, USA). Imaging was performed with ChemiDoc MP Imaging System (Bio-Rad, Hercules, CA, USA) with a 1 second exposure time. Imaging was also performed with a 13.5 second exposure time and a paper towel occluding the high molecular weight portion of the gel to gain better resolution of the short form RdmS bands in **Fig 4**. Coomassie blue dye (Thermo Fisher, Waltham, MA, USA) was used to evaluate near-equal loading of all cell lysates. All blots and Coomassie stained gel images are available in **Fig S8**.

### Readthrough efficiency calculation

We generated Eq. 1 by representing the rates of readthrough and truncation for the cellular proteome assuming second order kinetics as described in Eqs. 2 and 3. The rate of readthrough of any UAG codons in the cell depends on the concentration of Pyl-tRNA^Pyl^, the number of UAG-containing transcripts, and the rate constant k_readthrough_ as shown in Eq. 2. The rate of truncation of any UAG codon is a function of archaeal release factor (aRF1) concentration, the number of UAG-containing transcripts, and the rate constant k_truncation_ as shown in Eq. 3.

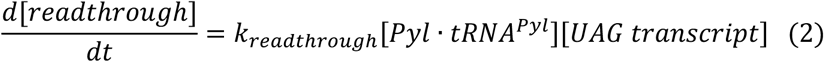

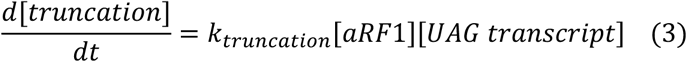

The rate of change of Pyl-tRNA^Pyl^ can be described by:

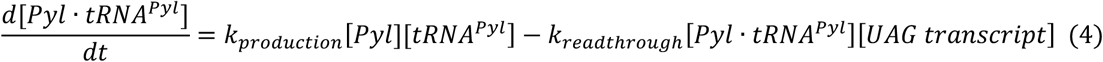

Where the first term on the right side of Eq. 4 represents the rate of production of Pyl-tRNA^Pyl^ and is dependent on the concentration of free pyrrolysine [Pyl], the uncharged tRNA^pyl^ [tRNA^pyl^] and the rate constant k_production_. The second term on the right represents the rate of consumption of Pyl-tRNA^Pyl^ due to readthrough of a UAG codon (i.e. the readthrough rate, Eq. 2). If we assume steady state concentrations of Pyl-tRNA^Pyl^ during growth on a given substrate then the production and consumption of Pyl-tRNA^Pyl^ are balanced, i.e.:

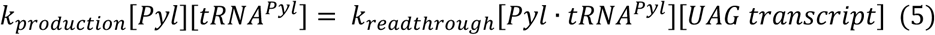

To determine the readthrough efficiency, we take the ratio of the readthrough rate—equivalent to the production rate in steady state (Eq. 5)—over the truncation rate (Eq. 3), yielding Eq. 6.

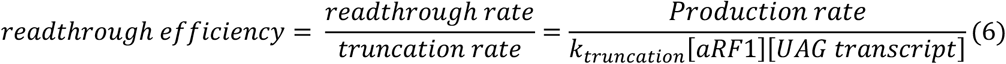

Since we observe that aRF1 transcription does not significantly change between MeOH and TMA growth conditions (**Fig S10C**, **Table S8**), we assume k_truncation_[aRF1] remains constant and find that:

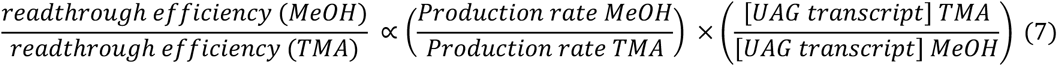

We refer to the Pyl-tRNA^Pyl^ production rate as the “Pyl supply” and estimate this value to be proportional to the FPKM (fragments per kilobase per million mapped reads) of the *pyl* genes for a given condition. We refer to the concentration of UAG transcripts as the “Pyl demand” and estimate it as the sum FPKM values for transcripts containing a UAG, using gene IDs identified in our Pyl ID script, for a given condition. This leads us from Eq. 7 to the relationship represented in Eq. 1 and illustrated in **Fig. 5**. This heuristic model for readthrough efficiency allows us to compare the ratio of Pyl-containing proteins to their truncated counterparts without quantifying the precise proportion of readthrough to truncation in any single condition. All FPKM values can be found in **Table S8**, and the calculations used to determine ratio of readthrough efficiencies between substrates can be found in **Table S10**.

## Data and code availability

Raw Illumina reads associated genome-wide DNA and RNA sequencing have been deposited to the Sequencing Reads Archive (SRA) and the project number will be made available upon publication. All custom scripts described can be found at the following repository: https://github.com/kshalv/pyl.

## Notes

### Competing Interest Statement

The authors have declared no competing interest.

### Summary of Updates

Citations edited in the main text

