## Supplementary Figures for "The Ambiguous Genetic Code of Methanogenic Archaea that Grow on Methylamines"

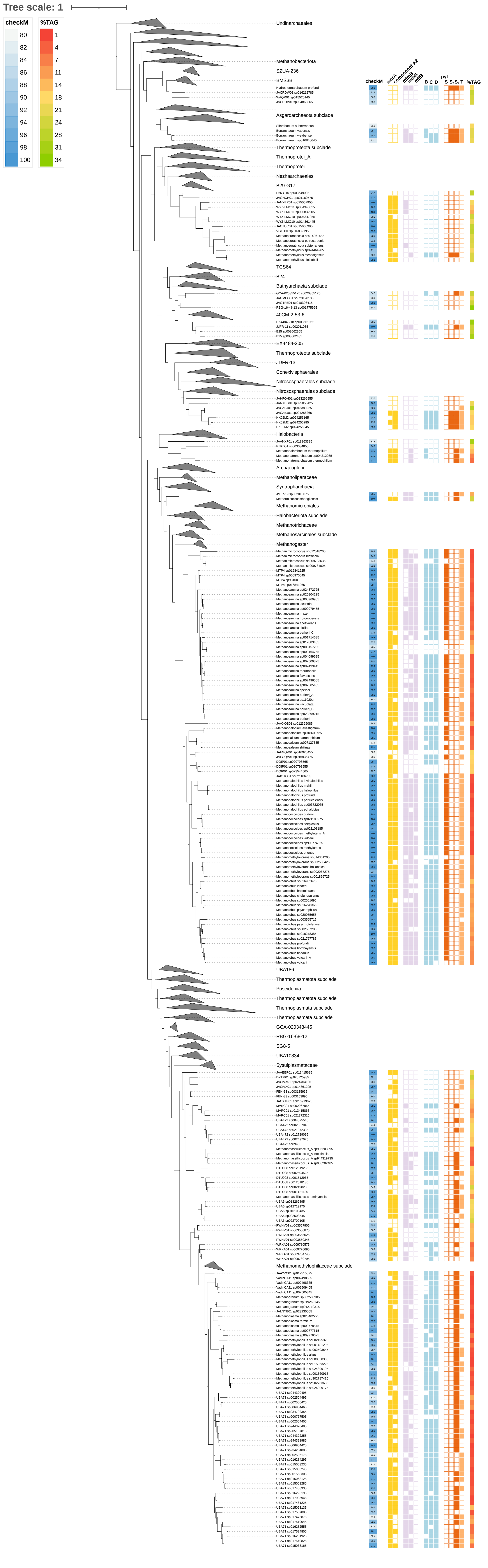
Tree scale: 1
checkM
80
82
84
86
88
90
92
94
96
98
100
%TAG
1
4
7
11
14
18
21
24
28
31
34
Undinarchaeales
Methanobacteriota
SZUA-236
BMS3B
Asgardarchaeota subclade
Thermoproteota subclade
Thermoprotei_A
Thermoprotei
Nezhaarchaeales
B29-G17
TCS64
B24
Bathyarchaeia subclade
40CM-2-53-6
EX4484-205
Thermoproteota subclade
JDFR-13
Conexivisphaerales
Nitrososphaerales subclade
Nitrososphaerales subclade
Halobacteria
Archaeoglobi
Methanoliparaceae
Syntropharchaeia
Methanomicrobiales
Halobacteriota subclade
Methanotrichaceae
Methanosarcinales subclade
Methanogaster
UBA186
Thermoplasmatota subclade
Poseidoniia
Thermoplasmatota subclade
Thermoplasmata subclade
Thermoplasmata subclade
GCA-020348445
RBG-16-68-12
SG8-5
UBA10834
Sysuiplasmataceae
Methanomethylophilaceae subclade
checkM
mcrA
component A2
mtmB
mtbB
mttB
B C D
pyl
S S₁ S₂ T
%TAG

**Fig S1.** Presence and absence of genes encoding the Pyrrolysine (Pyl) biosynthesis (*pylBCD*) and incorporation modules (*pylTS*), methylamine methyltransferases, methyl- coenzyme M reductase and *atwA* (component A2) (markers of archaeal methane metabolism) overlaid on the archaeal phylogeny from GTDB r214.0[1,2]. Genomes exceed a checkM completeness threshold of 80% and the completeness score is indicated in blue. TAG stop codon usage is indicated by the red-green gradient and clades universally lacking *pylBCD* and *pylTS* have been collapsed. **mcrA:** methyl coenzyme M reductase subunit Alpha; **mtmB:** mono-methylamine methyltransferase; **mtbB:** di-methylamine methyltransferase; **mttB:** tri-methylamine methyltransferase; **pylB:** methyl-ornithine synthase; **pylC:** 3-methylornithin L-lysine ligase; **pylD:** 3-methylornithyl-N6-L-lysine dehydrogenase; **pylS:** pyrrolysine $tRNA^{Pyl}$ ligase; **pylSn:** N-terminal subunit pyrrolysine $tRNA^{Pyl}$ ligase; **pylSc:** C-terminal subunit pyrrolysine $tRNA^{Pyl}$ ligase; **pylT:** $tRNA^{Pyl}$.

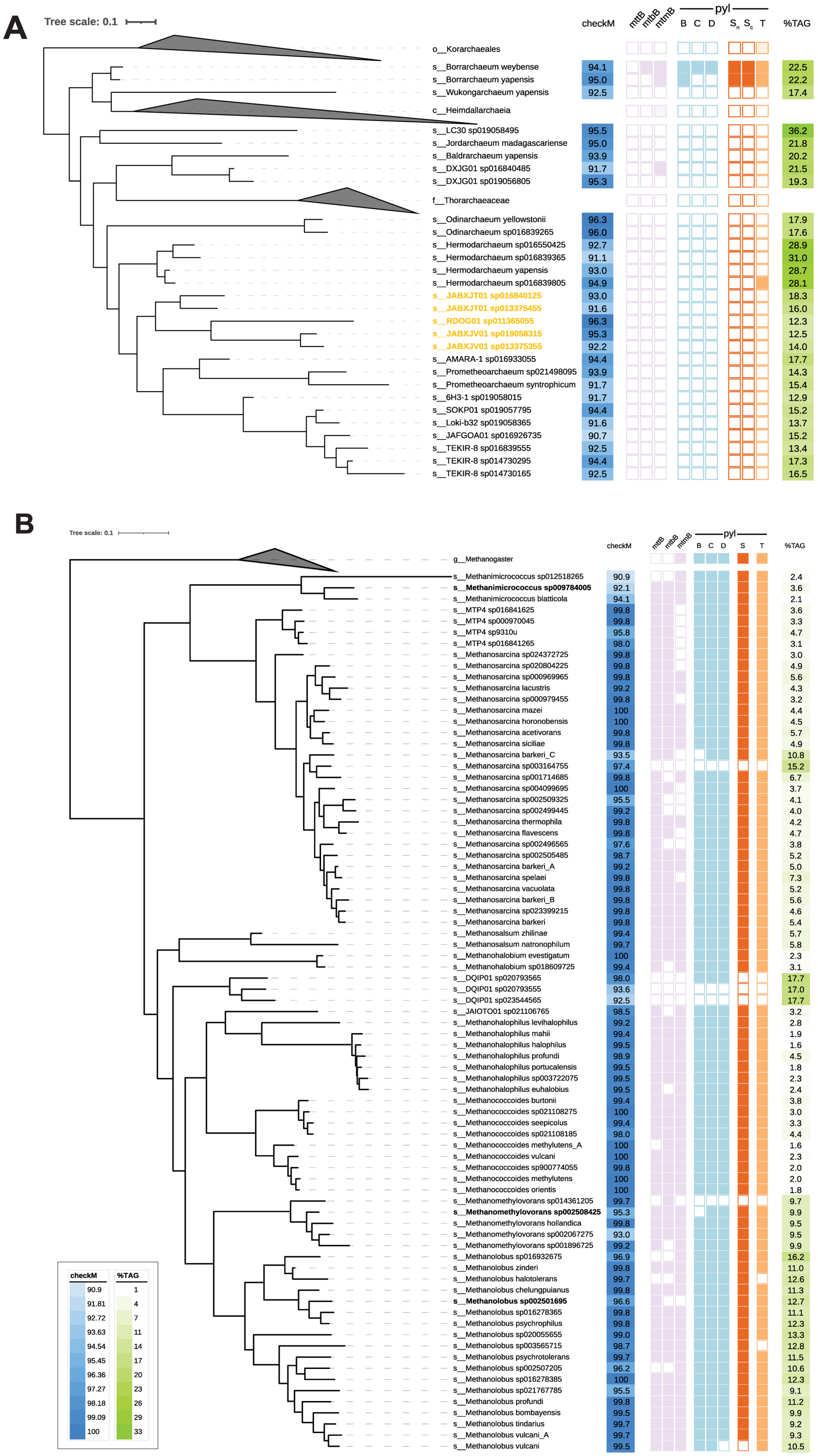


**Fig S2.** Pyl is found sporadically in *Asgardarchaeota* and widely in *Methanosarcinaceae*. All genomes exceed a CheckM threshold of 90%; scales are shared between trees. TAG stop codon usage is indicated by the white-green gradient. **A)** Tree for the phylum *Asgardarchaeota*; bolded yellow text indicates genomes that have the methane metabolism marker genes. **B)** Tree for the family *Methanosarcinaceae*; genomes in bold lack at least one of the following: *mcrA, mcrB, mcrG* or *atwA* (component A2). **mtbB:** di-methylamine methyltransferase; **mttB:** tri-methylamine methyltransferase; **pylB:** methyl-ornithine synthase; **pylC:** 3-methylornithine L-lysine ligase; **pylD:** 3-methylornithyl-N6-L-lysine dehydrogenase; **pylS:** pyrrolysine $tRNA^{Pyl}$ ligase; **pylSn:** N-terminal subunit pyrrolysine $tRNA^{Pyl}$ ligase; **pylSc:** C-terminal subunit pyrrolysine $tRNA^{Pyl}$ ligase; **pylT:** $tRNA^{Pyl}$.

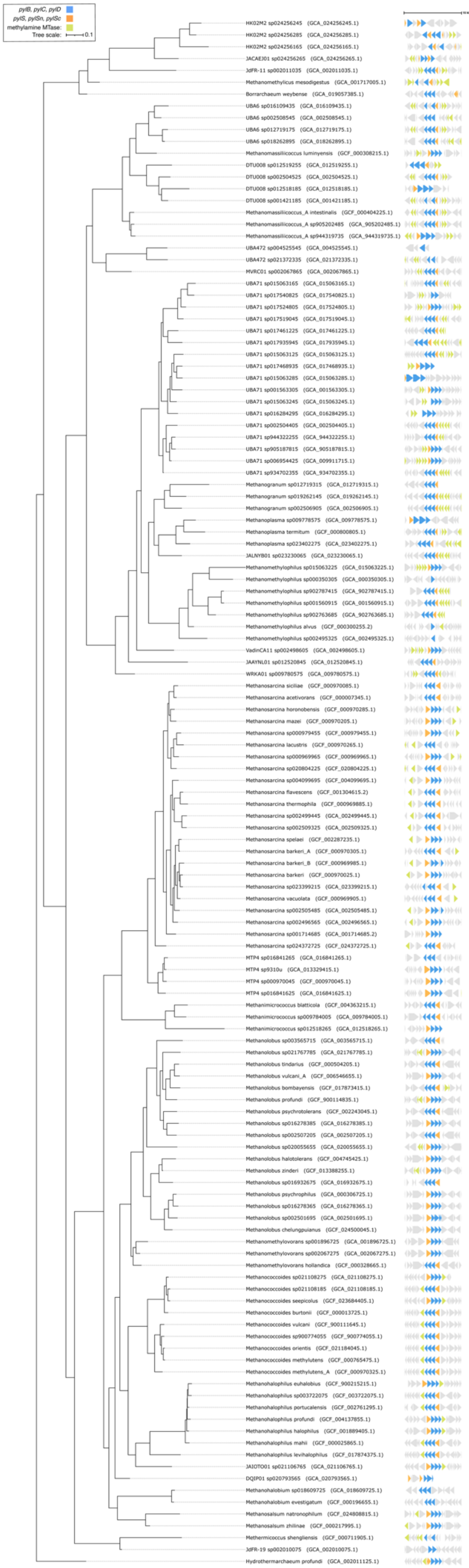

pylB, pylC, pylD
pylS, pylSn, pylSc
methylamine MTase:
Tree scale: 0.1
HK02M2 sp024256245 (GCA_024256245.1)
HK02M2 sp024256285 (GCA_024256285.1)
HK02M2 sp024256165 (GCA_024256165.1)
JACAEJ01 sp024256265 (GCA_024256265.1)
JdFR-11 sp002011035 (GCA_002011035.1)
Methanomethylicus mesodigestus (GCA_001717005.1)
Borrarchaeum weybense (GCA_019057385.1)
UBA6 sp016109435 (GCA_016109435.1)
UBA6 sp002508545 (GCA_002508545.1)
UBA6 sp012719175 (GCA_012719175.1)
UBA6 sp018262895 (GCA_018262895.1)
Methanomassiliicoccus luminyensis (GCF_000308215.1)
DTU008 sp012519255 (GCA_012519255.1)
DTU008 sp002504525 (GCA_002504525.1)
DTU008 sp012518185 (GCA_012518185.1)
DTU008 sp001421185 (GCA_001421185.1)
Methanomassiliicoccus_A intestinalis (GCF_000404225.1)
Methanomassiliicoccus_A sp905202485 (GCA_905202485.1)
Methanomassiliicoccus_A sp944319735 (GCA_944319735.1)
UBA472 sp004525545 (GCA_004525545.1)
UBA472 sp021372335 (GCA_021372335.1)
MVRC01 sp002067865 (GCA_002067865.1)
UBA71 sp015063165 (GCA_015063165.1)
UBA71 sp017540825 (GCA_017540825.1)
UBA71 sp017524805 (GCA_017524805.1)
UBA71 sp017519045 (GCA_017519045.1)
UBA71 sp017461225 (GCA_017461225.1)
UBA71 sp017935945 (GCA_017935945.1)
UBA71 sp015063125 (GCA_015063125.1)
UBA71 sp017468935 (GCA_017468935.1)
UBA71 sp015063285 (GCA_015063285.1)
UBA71 sp001563305 (GCA_001563305.1)
UBA71 sp015063245 (GCA_015063245.1)
UBA71 sp016284295 (GCA_016284295.1)
UBA71 sp002504405 (GCA_002504405.1)
UBA71 sp944322255 (GCA_944322255.1)
UBA71 sp905187815 (GCA_905187815.1)
UBA71 sp006954425 (GCA_009911715.1)
UBA71 sp934702355 (GCA_934702355.1)
Methanogranum sp012719315 (GCA_012719315.1)
Methanogranum sp019262145 (GCA_019262145.1)
Methanogranum sp002506905 (GCA_002506905.1)
Methanoplasma sp009778575 (GCA_009778575.1)
Methanoplasma termitum (GCF_000800805.1)
Methanoplasma sp023402275 (GCA_023402275.1)
JALNYB01 sp023230065 (GCA_023230065.1)
Methanomethylophilus sp015063225 (GCA_015063225.1)
Methanomethylophilus sp000350305 (GCA_000350305.1)
Methanomethylophilus sp902787415 (GCA_902787415.1)
Methanomethylophilus sp001560915 (GCA_001560915.1)
Methanomethylophilus sp902763685 (GCA_902763685.1)
Methanomethylophilus alvus (GCF_000300255.2)
Methanomethylophilus sp002495325 (GCA_002495325.1)
VadinCA11 sp002498605 (GCA_002498605.1)
JAAYNL01 sp012520845 (GCA_012520845.1)
WRKA01 sp009780575 (GCA_009780575.1)
Methanosarcina siciliae (GCF_000970085.1)
Methanosarcina acetivorans (GCF_000007345.1)
Methanosarcina horonobensis (GCF_000970285.1)
Methanosarcina mazei (GCF_000970205.1)
Methanosarcina sp000979455 (GCF_000979455.1)
Methanosarcina lacustris (GCF_000970265.1)
Methanosarcina sp000969965 (GCF_000969965.1)
Methanosarcina sp020804225 (GCF_020804225.1)
Methanosarcina sp004099695 (GCF_004099695.1)
Methanosarcina flavescens (GCF_001304615.2)
Methanosarcina thermophila (GCF_000969885.1)
Methanosarcina sp002499445 (GCA_002499445.1)
Methanosarcina sp002509325 (GCA_002509325.1)
Methanosarcina spelaei (GCF_002287235.1)
Methanosarcina barkeri_A (GCF_000970305.1)
Methanosarcina barkeri_B (GCF_000969985.1)
Methanosarcina barkeri (GCF_000970025.1)
Methanosarcina sp023399215 (GCA_023399215.1)
Methanosarcina vacuolata (GCF_000969905.1)
Methanosarcina sp002505485 (GCA_002505485.1)
Methanosarcina sp002496565 (GCA_002496565.1)
Methanosarcina sp001714685 (GCA_001714685.2)
Methanosarcina sp024372725 (GCF_024372725.1)
MTP4 sp016841265 (GCA_016841265.1)
MTP4 sp9310u (GCA_013329415.1)
MTP4 sp000970045 (GCF_000970045.1)
MTP4 sp016841625 (GCA_016841625.1)
Methanimicrococcus blatticola (GCF_004363215.1)
Methanimicrococcus sp009784005 (GCA_009784005.1)
Methanimicrococcus sp012518265 (GCA_012518265.1)
Methanolobus sp003565715 (GCA_003565715.1)
Methanolobus sp021767785 (GCA_021767785.1)
Methanolobus tindarius (GCF_000504205.1)
Methanolobus vulcani_A (GCF_006546655.1)
Methanolobus bombayensis (GCF_017873415.1)
Methanolobus profundi (GCF_900114835.1)
Methanolobus psychrotolerans (GCF_002243045.1)
Methanolobus sp016278385 (GCA_016278385.1)
Methanolobus sp002507205 (GCA_002507205.1)
Methanolobus sp020055655 (GCA_020055655.1)
Methanolobus halotolerans (GCF_004745425.1)
Methanolobus zinderi (GCF_013388255.1)
Methanolobus sp016932675 (GCA_016932675.1)
Methanolobus psychrophilus (GCA_000306725.1)
Methanolobus sp016278365 (GCA_016278365.1)
Methanolobus sp002501695 (GCA_002501695.1)
Methanolobus chelungpuianus (GCF_024500045.1)
Methanomethylovorans sp001896725 (GCA_001896725.1)
Methanomethylovorans sp002067275 (GCA_002067275.1)
Methanomethylovorans hollandica (GCF_000328665.1)
Methanococcoides sp021108275 (GCA_021108275.1)
Methanococcoides sp021108185 (GCA_021108185.1)
Methanococcoides seepicolus (GCF_023684405.1)
Methanococcoides burtonii (GCF_000013725.1)
Methanococcoides vulcani (GCF_900111645.1)
Methanococcoides sp900774055 (GCF_900774055.1)
Methanococcoides orientis (GCF_021184045.1)
Methanococcoides methylutens (GCF_000765475.1)
Methanococcoides methylutens_A (GCF_000970325.1)
Methanohalophilus euhalobius (GCF_900215215.1)
Methanohalophilus sp003722075 (GCF_003722075.1)
Methanohalophilus portucalensis (GCF_002761295.1)
Methanohalophilus profundi (GCF_004137855.1)
Methanohalophilus halophilus (GCF_001889405.1)
Methanohalophilus mahii (GCF_000025865.1)
Methanohalophilus levihalophilus (GCF_017874375.1)
JAIOTO01 sp021106765 (GCA_021106765.1)
DQIP01 sp020793565 (GCA_020793565.1)
Methanohalobium sp018609725 (GCA_018609725.1)
Methanohalobium evestigatum (GCF_000196655.1)
Methanosalsum natronophilum (GCF_024808815.1)
Methanosalsum zhilinae (GCF_000217995.1)
Methermicoccus shengliensis (GCF_000711905.1)
JdFR-19 sp002010075 (GCA_002010075.1)
Hydrothermarchaeum profundi (GCA_002011125.1)

**Fig S3.** Genomic neighborhood of Pyrrolysine (Pyl) biosynthetic machinery in $Pyl^+$ genomes (i.e. genomes that encode *pylBCD, pylT,* and *pylRS*). Pyl biosynthetic machinery is in blue, Pyl incorporation genes are in orange, and methylamine-specific methyltransferases are in green. Genome neighborhoods are centered around *pylB* and include 7.5 kb up and downstream of the start and stop sites. **MTase:** methyltransferase; **pylB:** methyl-ornithine synthase; **pylC:** 3-methylornithin L-lysine ligase; **pylD:** 3-methylornithyl-N6-L-lysine dehydrogenase; **pylS:** pyrrolysine $tRNA^{Pyl}$ ligase; **pylSn:** N-terminal subunit pyrrolysine $tRNA^{Pyl}$ ligase; **pylSc:** C-terminal subunit pyrrolysine $tRNA^{Pyl}$ ligase; **pylT:** $tRNA^{Pyl}$.

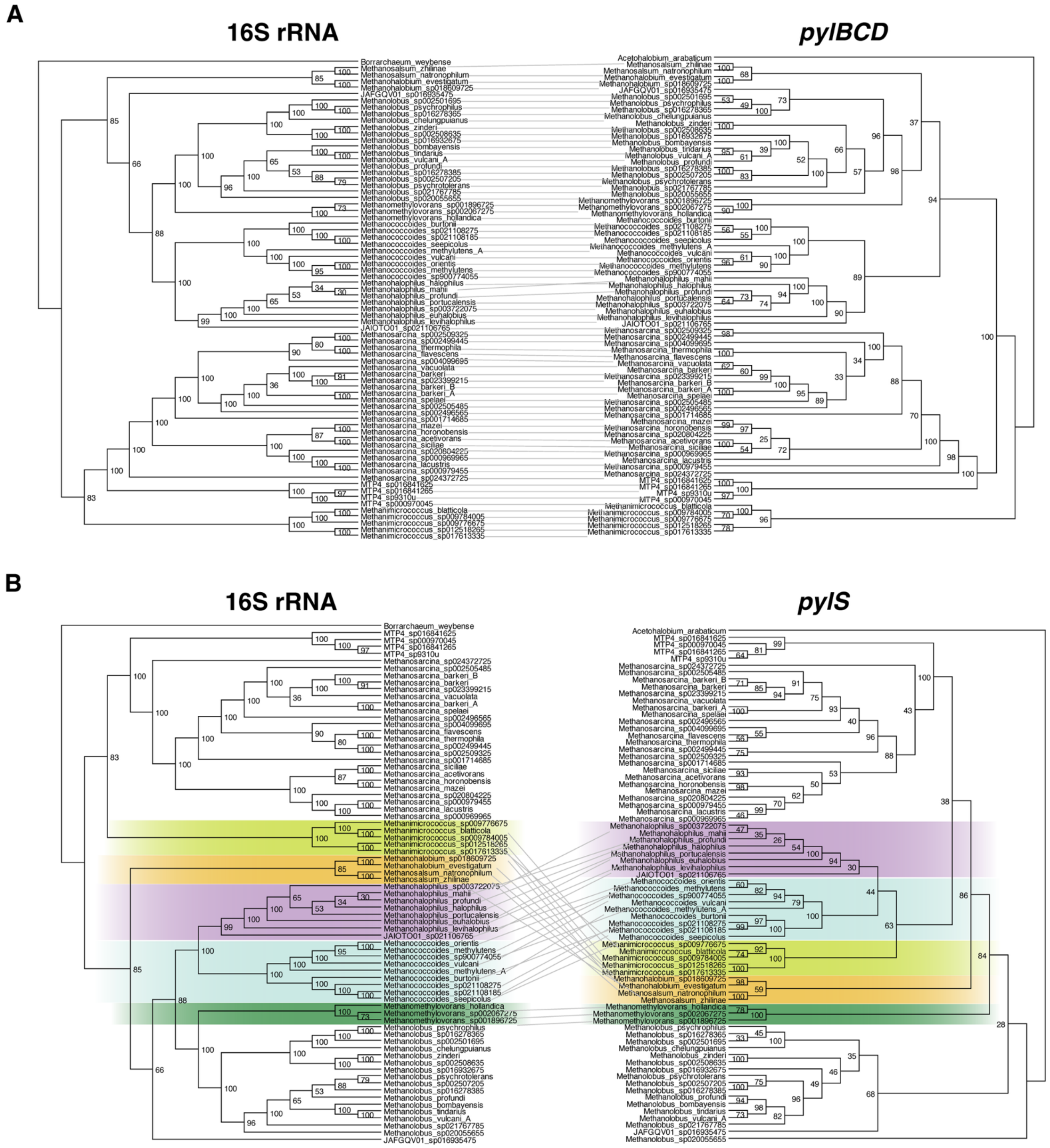


**Fig S4.** Tanglegram comparing the gene trees for the Pyrrolysine (Pyl) biosynthetic and incorporation components and the 16S rRNA tree for members of the family *Methanosarcinaceae.* All 16S ribosomal RNA trees are rooted to the 16S rRNA of *Borrarchaeum weybense*, and gene trees are rooted to the bacterial Pyl sequences from *Acetohalobium arabaticum*. A tanglegram for **A)** concatenated *pylBCD* alignments and **B)** *pylS* alignments, contrasted with the species tree of the family *Methanosarcinaceae.* Notable differences in tree topologies are highlighted in colors distinguished by genus taxonomy. Species tree was generated by RAxML using an alignment of 16S rRNA gene sequences derived from GTDB r214.0. Gene trees were generated using a MAFFT alignment, which was trimmed at 90% with TrimAl, concatenated, and assembled with RAxML 8.2.11 plug-in (GAMMA BLOSUM62 matrix, rapid-hill climbing with bootstrapping, n=100). Node labels represent bootstrap values.

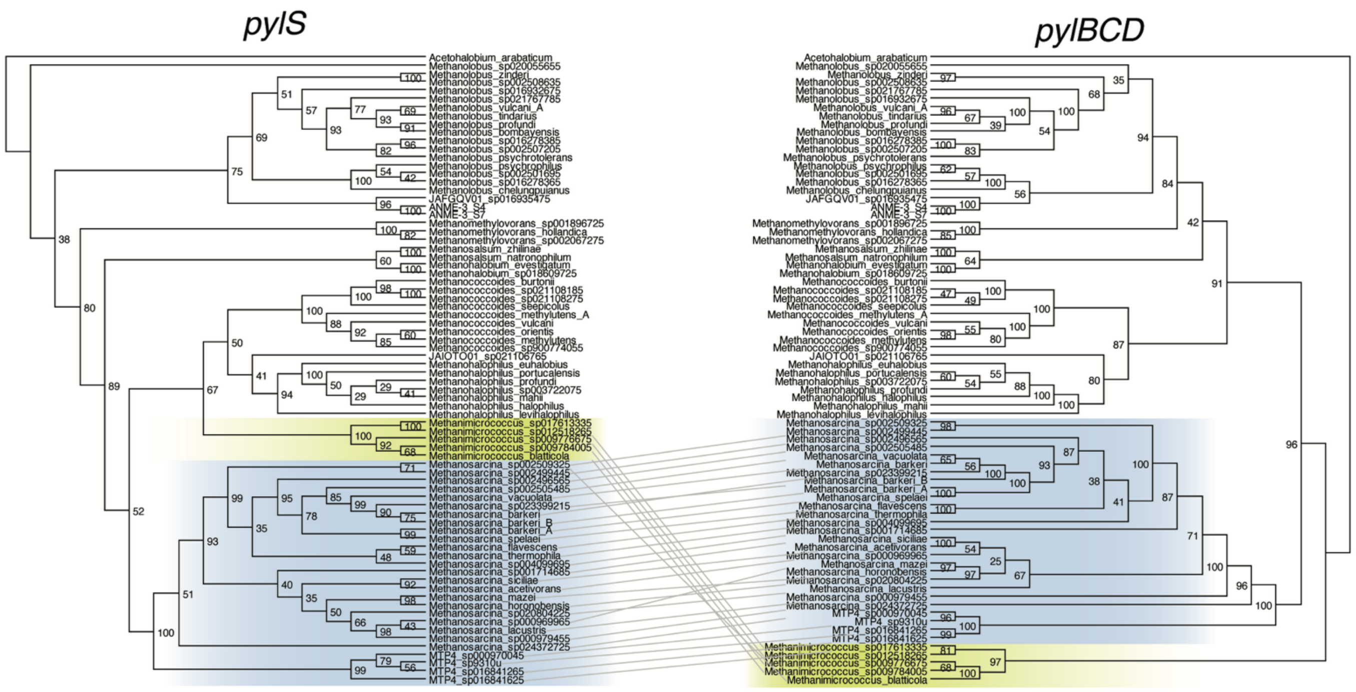

**Fig S5.** A tanglegram contrasting the Pyl incorporation (*pylS*) and biosynthetic (*pylBCD*) components from Fig S4. Both trees are rooted to the bacterial Pyl+ genome, *Acetohalobium arabaticum*. Significant differences in tree topology are highlighted by genus in color. Tree was generated with RAxML 8.2.11 (GAMMA BLOSUM62, n =100). Node labels represent bootstrap values.

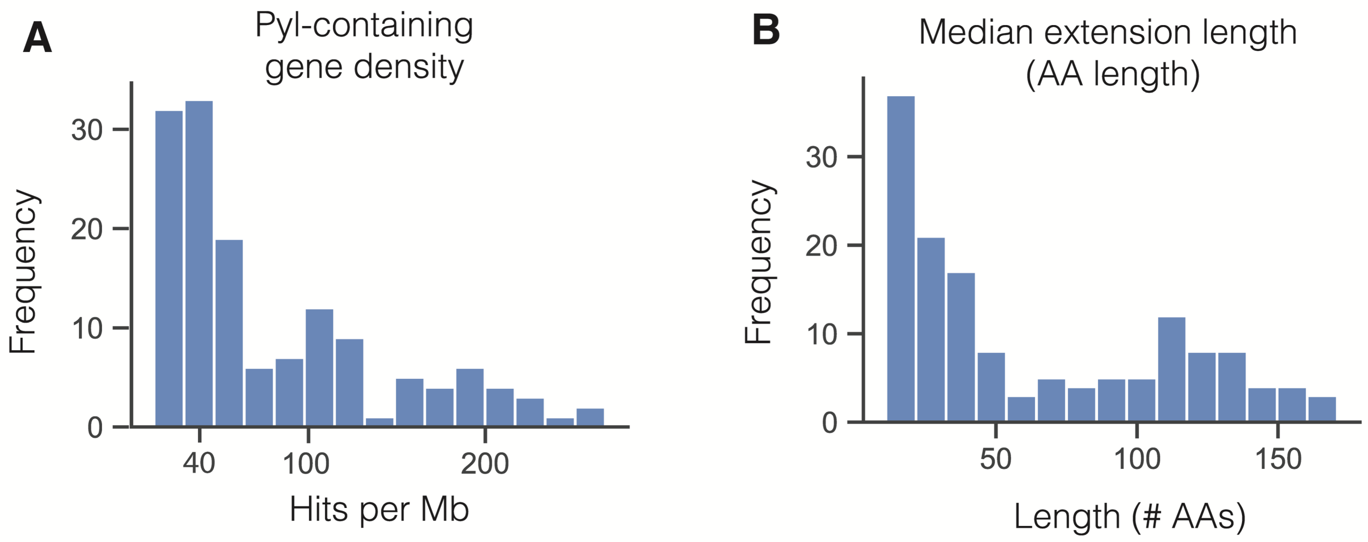


**Fig S6.** Histograms depicting genomic features of predicted Pyrrolysine (Pyl) containing genes. **A)** Pyl-containing gene density. Per genome, putative Pyl-containing genes were counted and normalized to genome size (Mb). **B)** Median extension length for predicted Pyl-containing genes per genome. Extension lengths were calculated as the distance between the first Pyl residue in an open reading frame and a downstream alternate stop codon (TAA or TGA).

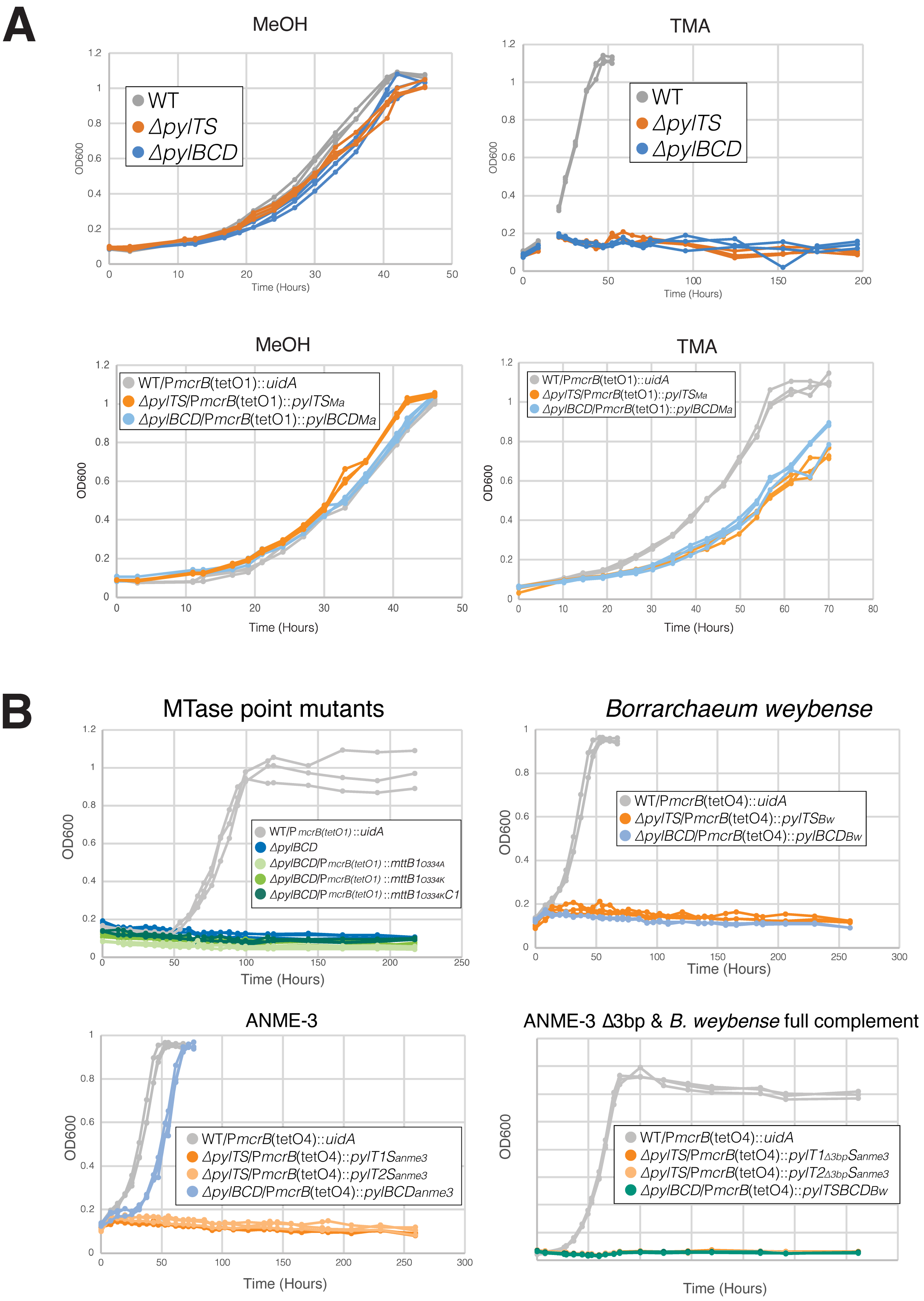


**Fig S7.** Triplicate growth curves for experiments using *Methanosarcina acetivorans* as a heterologous host to express *pyl* genes from archaea. **A)** Growth curves of WWM60 (denoted as wildtype or WT; in gray), single deletion knockouts of the Pyl biosynthetic (Δ*pylBCD;* blue) and incorporation genes (Δ*pylTS;* orange) in the WWM60 background in high-salt (HS) medium supplemented with either 125 mM methanol (MeOH) or 50 mM trimethylamine (TMA) as indicated. All complementation mutants express the deleted genes *in trans* from a tetracycline inducible promoter P*mcrB*(*tetO1*). For WT, a control gene, encoding β-glucuronidase*, uidA*, was expressed *in trans* on a plasmid. Growth curves were conducted in medium supplemented with puromycin at 2 µg/ml (to maintain the plasmid) and tetracycline at 100 µg/ml(to fully induce gene expression). **B)** Growth curves for TMA methyltransferase (*mttB*) point mutants and all heterologous expression mutants. The *mttB1* point mutants include: $mttB_{O334A}$ (light green), $mttB_{O334K}$ (green), and $mttB_{O334K}C$ (dark green). Heterologous expression strains include: $pylTS_{Bw}$ (orange), $pylT1S_{anme\text{-}3}$ (dark orange), $pylT2S_{anme\text{-}3}$ (light orange), $pylBCD_{Bw}$ (blue), $pylBCD_{anme\text{-}3}$ (blue), $pylT1_{\Delta 3bp}S_{anme\text{-}3}$ (dark orange), $pylT2_{\Delta 3bp}S_{anme\text{-}3}$ (light orange), and $pylTSBCD_{Bw}$ (teal). All genes were expressed from a tetracycline inducible promoter P*mcrB*(*tetO1*) (*mttB* point mutants) or at P*mcrB*(tetO4) (non-native Pyl machinery) *in* trans on a plasmid. Growth curves were conducted in medium supplemented with puromycin at 2 µg/ml (to maintain the plasmid) and tetracycline at 100 µg/ml(to fully induce gene expression). Each point mutation is indicated in the graphical legend using the following amino acid codes: **O:** Pyrrolysine; **A:** Alanine; **K:** Lysine. ***Ma:*** *Methanosarcina acetivorans;* ***Bw:*** *Borrarchaeum weybense.*

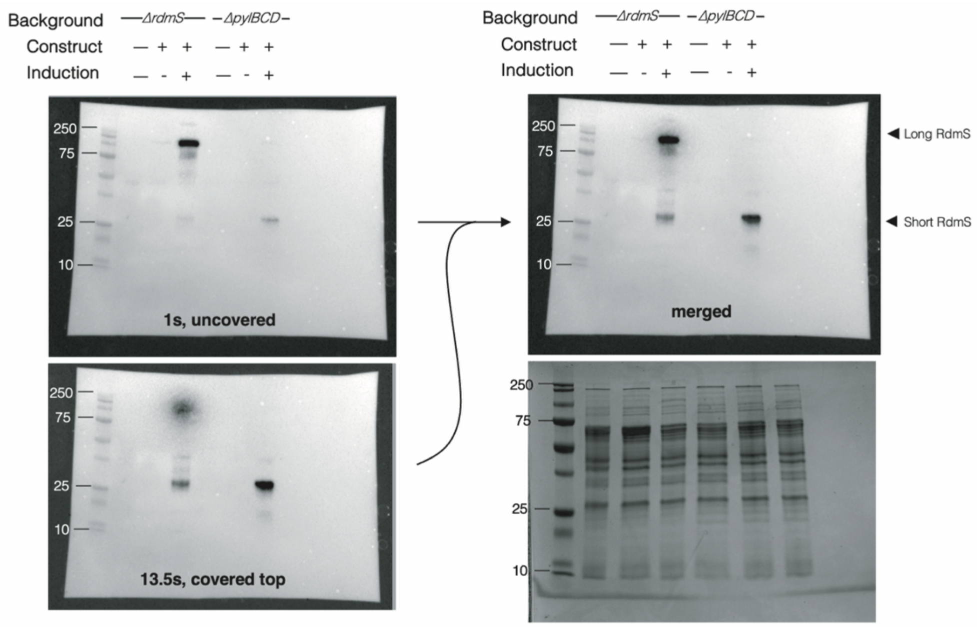


**Fig S8.** Full images for the anti-FLAG Western blot and the Coomassie stained protein gels shown in Fig 4A. The membrane was imaged twice (left) to gain a higher resolution image of the short-form RdmS band. The membrane was first exposed for 1s (top left), then, upper bands were covered to prevent oversaturation, and the membrane was imaged for 13.5s (bottom left). The two images were then merged (top right). Coomassie stained SDS-PAGE gel is shown as a loading control (bottom right), for near equal loading of 30 µg crude cell lysates. Plus (+) symbols indicate the presence of a plasmid construct expressing the target gene from a tetracycline-inducible promoter (P*mcrB*(*tetO4*)) at full induction (100 µg/ml tetracycline) with puromycin (2 µg/ml) added to maintain the plasmid.

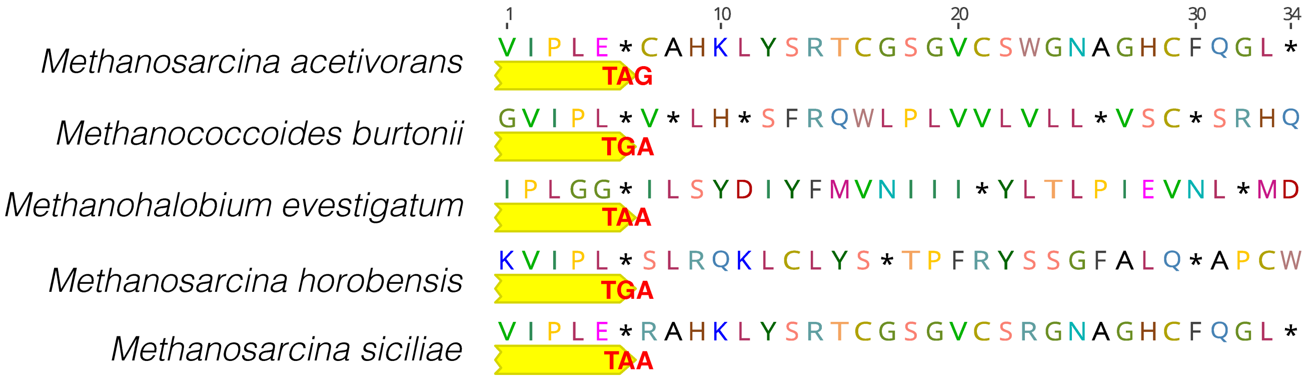


**Fig S9.** RnfA amino acid sequences from members of *Methanosarcinaceae* with different stop codon identities. Sequences shown are from the region five amino acids upstream of the stop codon to 28 amino acids downstream of the stop codon, which corresponds with the predicted 28 AA extension for *rnfA* from *Methanosarcina acetivorans*. The stop codon identity is indicated in bolded red text on the gene annotation beneath the amino acid sequence. Annotations and translations for the downstream *rnfB* sequence were excluded for clarity.

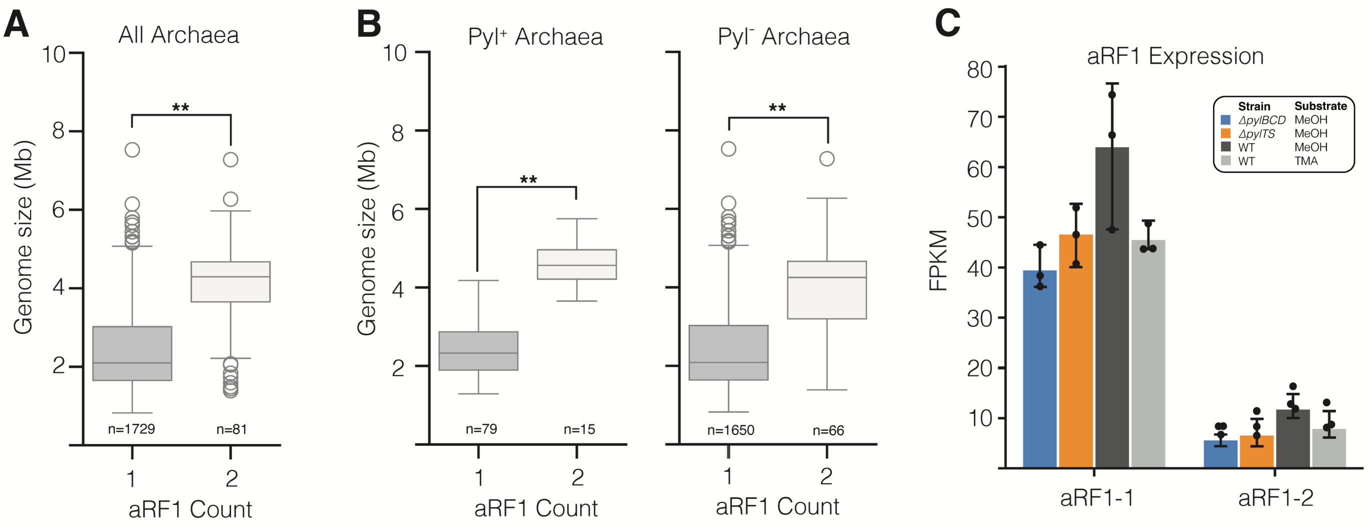


**Fig S10.** Box and whisker plots describing the relationship between archaeal release factor (aRF1) copy number and genome size. **A)** Relationship between aRF1 copy number and genome size across all archaea. **B)** Relationship between aRF1 and genome size within Pyl$^+$ and Pyl$^-$ genomes within archaea. **C)** FPKM (Fragment per kilobase per million reads) values for aRF1-1 and aRF1-2 in the Δ*pylBCD* mutant on methanol (MeOH) (blue), Δ*pylTS* mutant on MeOH (blue), WWM60 (parent strain also referred to as wildtype or WT) on MeOH (light gray) or trimethylamine (TMA) (dark gray). Error bars represent the standard deviation in triplicate FPKM values. No pairwise comparison of FPKM values for aRF1 copies by DESeq2 were called as significantly differentially expressed ($q<0.001$ and $\log_2$-fold≤-1.5 or $\log_2$-fold≥1.5); full data can be found in Table S8. All statistical tests in panels A and B were conducted using Welch's t-test where $^{**}p<0.001$. Note that we only consider hits for aRF1, and not eRF1 in this analysis given that its function in archaea is not yet mapped.
